# A quantitative approach to poxvirus infection reveals general correlates of antigen presentation and immunogenicity for viral CD8^+^ T cell epitopes

**DOI:** 10.1101/2024.11.18.624047

**Authors:** Matthew J Witney, Nathan P Croft, Yik Chun Wong, Stewart A Smith, Inge EA Flesch, Erica Keller, Nicole L La Gruta, Anthony W Purcell, David C. Tscharke

**Author notes:** Co-first authors with equal contribution (Matthew J Witney, Nathan P Croft). Co-senior authors with equal contribution (Anthony W Purcell, David C. Tscharke).

## Abstract

CD8^+^ T cells are essential effectors in antiviral immunity that kill infected cells displaying viral peptide epitopes on MHC class I. The pathways underpinning antigen presentation on MHC I are well known but we lack a general understanding of the quantitative and kinetic relationships between source proteins and presented epitopes, and how these relate to immunogenicity. We used quantitative mass spectrometry to interrogate infection with vaccinia virus, the MPOX vaccine, measuring amounts of >40 viral epitopes and their source proteins multiple times after infection in four cell types. This revealed that up to 90% of the epitopes were presented as fast as their source proteins were made and that protein amounts failed to correlate with epitope levels. Further, a significant correlation between epitope levels and immunogenicity required presentation to be quantified *ex vivo*, with each step away from a matched infection route and natural presenting cell eroding the association.

## Introduction

CD8^+^ T cells are critical effectors during viral infection that detect and eradicate infected cells presenting viral peptides on MHC Class I. Peptides presented on MHC Class I (p:MHC-I) are typically short 8-16mers, derived from proteins degraded and processed within the cell ^1, 2^. During virus infection, p:MHC-I derived from viral proteins can be recognized by T cell receptors (TCR)s on CD8^+^ T cells. Typically, the majority of the anti-viral CD8^+^ T cell response is biased towards a relatively small number of viral epitopes. Further, the size of response differs across these epitopes in a reproducible manner that can span two orders of magnitude, allowing a ranking of epitopes by immunogenicity referred to as the immunodominance hierarchy ^3–5^.

The source of peptides presented on MHC-I on most cell types is from endogenously translated protein antigens. These are degraded predominantly by proteasomes and the fragments are shuttled into the endoplasmic reticulum and loaded onto MHC-I by a complex that includes a specialised transporter and chaperones. Once assembled, new p:MHC-Is can be subject to further trimming of the peptide and selection for adequate stability before being exported to the cell surface. However, not all cell types express the same antigen processing machinery, potentially altering how a particular antigen is processed and presented. Most notably this includes the expression of three alternative subunits of the proteasome (LMP2, LMP7 and MECL-1), which replace catalytically active subunits of the proteasome to form the immunoproteasome ^6–11^ and exhibit biased proteolytic degradation of proteins compared to constitutive proteasomes ^12–14^. These subunits are constitutively expressed in professional antigen presenting cells (APCs), such as dendritic cells (DCs) and can also be induced by inflammatory cytokines in other cell types.

At the start of an infection, naïve CD8^+^ T cells must engage a cognate p:MHC-I with their TCR on activated DCs to initiate a program of clonal expansion and differentiation into mature, or primed effectors that can then kill infected cells displaying the same p:MHC-I. This obligatory role of activated DCs ensures that CD8^+^ T cells are only deployed in the context of infection or other physiological threats. Where a particular virus can infect DCs, the endogenous antigen presentation pathway as described above is adequate to prime a CD8^+^ T cell response. However, DCs are uniquely able to take up antigen from other cells or the extracellular space and process these for presentation on MHC-I. This is referred to as cross-presentation and allows the priming of CD8^+^ T cells irrespective of the infection of DCs ^15^.

Although the individual components of the antigen processing and presentation machinery are well described (reviewed by Pishesha, Harmand ^2^), the primary source of most epitopes, the relationship between antigen abundance and epitope levels, and the extent to which these might differ across cell types remain contentious or lacks a broader context that accounts for diversity in the proteome and immunopeptidome. In particular, while it is accepted that peptides may be derived from degradation of early translational products, such as defective ribosomal products (DRiPs) ^16–19^, or from the steady-state turnover of metabolically mature proteins (so called ‘retirees’ ^20^), the fraction of presented peptides during viral infection derived from either source remains controversial ^21, 22^. Similarly, although peptides presented on MHC-I derived from abundant proteins are more likely to be identified by mass spectrometry ^23^, whether the abundance of the source protein is a general correlate of epitope presentation levels is not clear, particularly for virus-derived antigens ^24^. Finally, while some studies have found that presentation differs markedly between two cell types for selected epitopes with evidence of biological relevance, without data on a wide range of epitopes it is unclear whether these differences are outliers or the norm ^25, 26^.

Understanding the factors that contribute to epitope abundance is viewed to be important because it is widely assumed to be a general correlate of the number of peptide-specific CD8^+^ T cell responses during viral infection ^5^. The largest body of data underpinning this idea are from indirect measures related to presentation, such as p:MHC-I affinity and stability, that are considered to reflect epitope levels ^27^. However, there is disagreement across studies about the limits of whether p:MHC-I predicts the magnitude of responses or is better thought of as a parameter that helps discriminate immunogenic from non-immunogenic peptides ^24, 28, 29^. The other strand of evidence is from studies showing that increasing the amount of a single model peptide presented increases the size of CD8^+^ T cell responses ^30, 31^. Approaches that have directly quantified natural epitope abundance frequently investigate too few epitopes to allow for statistically meaningful conclusions ^32–39^. Moreover, how quantitative features of epitopes are affected by the choice of cell type *in vitro,* and the extent this reflects presentation *in vivo* remains entirely unexplored.

In the largest study to investigate quantitative features of the virus-derived immunopeptidome to date, the abundance of 22 epitopes, source antigen and the peptide-specific CD8^+^ T cells were measured in the context of influenza A infection in mice ^24^. This work found that epitope levels were disconnected from the abundance of the source antigen, but a statistically significant correlation between the abundance of 22 epitopes and peptide-specific CD8^+^ T cell responses was identified ^24^. It is unclear how well these findings can be generalized, in part because around 85% of all CD8^+^ T cells in the response to this virus in mice can be accounted for by only six peptides, with two peptides contributing around two thirds of the total response, reducing actual complexity. Further, a single dominant peptide changed the best correlate of immunogenicity from presentation on infected cells to presentation on cross presenting DCs. The next largest study quantified the abundance of nine p:MHC-I on vaccinia virus (VACV) infected cells and failed to identify a correlation between levels of these epitopes and CD8^+^ T cell responses ^40^. This study employed less rigorous quantification, and all peptides were presented by a single MHC-I allomorph (H-2K^b^), leaving the question of presentation levels and immunogenicity unanswered.

To answer these questions definitively, what is required is a viral infection model in which there are a large number of well-defined epitopes that evenly span a wide range of abundance and to which CD8^+^ T cell responses are also well spread. VACV has long been used as a tractable model pathogen for studying CD8^+^ T cell responses, having a large and complex proteome and robust innate and adaptive immune responses during infection ^41^. VACV has 170 mass spectrometry-validated p:MHC-I for which immunogenicity has been rigorously tested ^4, 40^ and mouse models of infection are strictly acute with direct presentation of epitope on infected DC directly priming CD8+ T cells ^42–48^. Finally, VACV has current relevance as the type species of the orthopoxvirus family and the vaccine for MPOX.

Here we have extended the VACV model, collecting large mass spectrometry datasets from four cell types infected *in vitro* to model primary and immortalised fibroblasts and DCs. This information was used to comprehensively address how source protein expression kinetics and abundance relate to epitope presentation. We then addressed the relationship between epitope abundance and immunogenicity: first using our *in vitro* epitope presentation data and then in a new model in which we extended the current state-of-the-art in mass spectrometry to measure the abundance of epitope presentation *ex vivo* from uninfected mice. All sources of epitope and T cell information were considered to find that epitope presentation levels are indeed correlated with the magnitude of CD8^+^ T cell responses, but demonstrating this relationship requires a close matching of the models used to measure epitopes and T cells. These are the largest mass spectrometry datasets for a poxvirus, or indeed any virus infection. Our analysis considers multiple infection models, providing confidence that the findings here will be broadly generalizable across virus infections and, most likely, antigen presentation as a whole.

## Results

### Abundance of 45 VACV p:MHC-I on cells infected *in vitro*

To measure the abundance of 45 p:MHC-I on VACV-infected cell types, we used the sensitive mass spectrometry technique of Multiple Reaction Monitoring (MRM) MS, using internal isotopically labelled standard peptides to provide accurate quantification of endogenously presented viral peptides (Fig. 1A). Cells were infected *in vitro* with VACV between 0.5 and 8.5h to allow time for the complete VACV replication cycle ^16^. The absolute abundance of peptide detected by mass spectrometry can be normalized to the number of cells, providing a measure of epitope density per cell (Fig. 1B). Using this strategy, the abundance of 45 VACV epitopes was simultaneously measured on different antigen presenting cells, specifically bone-marrow derived dendritic cells (BMDCs), primary murine fibroblasts (PMFs), as well as the dendritic-like and fibroblastic-like cell lines, DC2.4 and MC57G, respectively ^49, 50^ (Fig. 1C-D).

**Figure 1:**
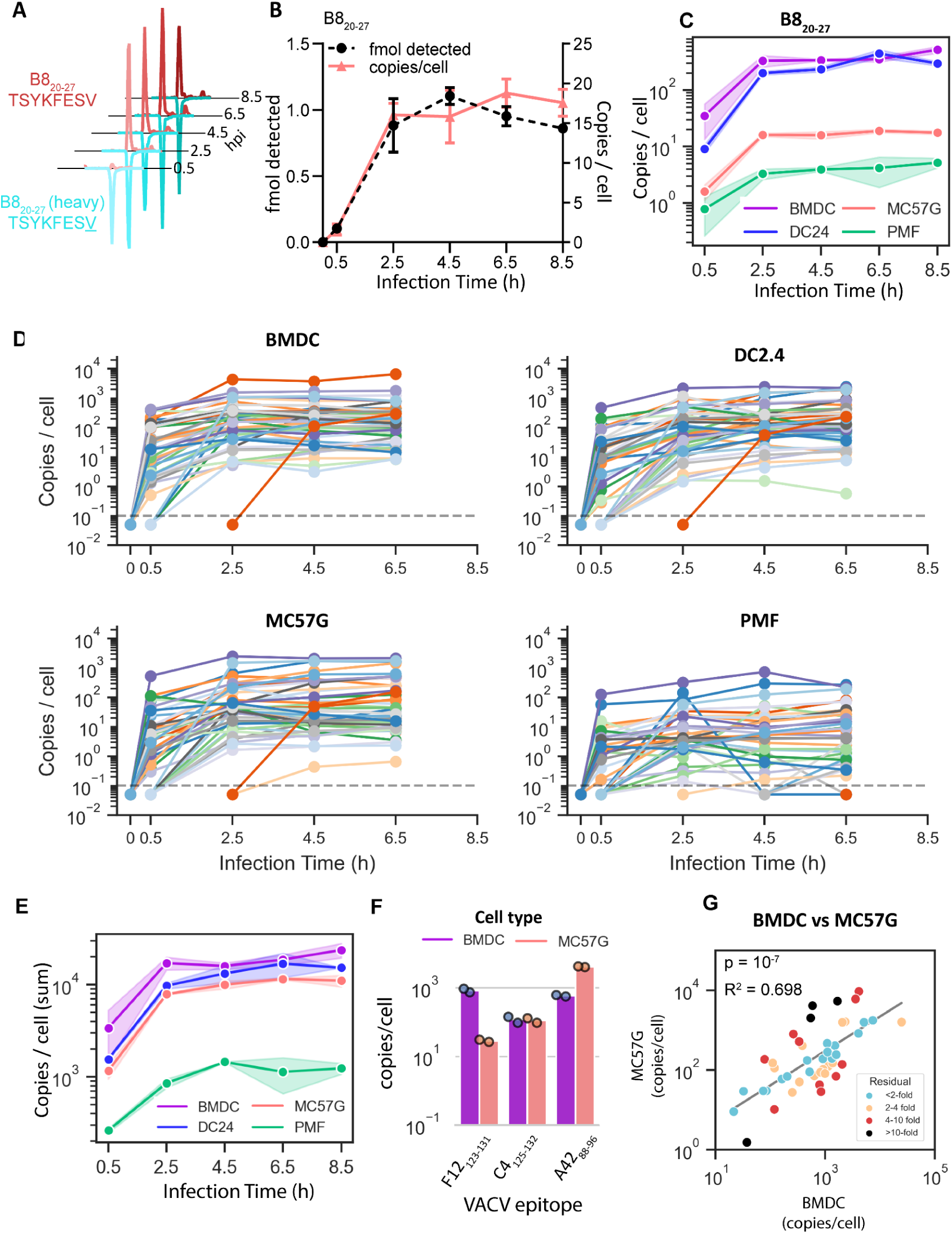
Quantification of 45 VACV epitope presentation on infected cells over time. Cells were infected *in vitro* with VACV for 0.5 h and up to 8.5 hr. Epitope presentation was quantified by MRM MS. A) Chromatogram peaks representing detection of VACV-specific and isotopically labelled B8_20-27_ peptide presented on VACV-infected MC57G cells at timepoints between 0.5h and 8.5h p.i. B) B8_20-27_ peptide detected by mass spectrometry (black), or calculated B8_20-27_ peptide copies per cell (pink), at each timepoint on MC57G cells. C) Abundance of B8_20-27_ presented on H-2K^b^ over time in BMDC, DC2.4, MC57G and PMF cells. Shaded regions represent the range from two independent replicates. D) Abundance of 45 VACV epitopes over time presented on BMDC, DC2.4, MC57G and PMF cells. Lines represent the average abundance of an epitope from two independent replicates and are drawn from the timepoint prior to time of first detection. Points below the dotted grey line identify undetected peptides. E) The sum abundance of all 45 epitopes on each cell type at each timepoint. Shaded regions represent the range from two independent replicates. F) The sum of three epitope abundances over time for three VACV epitopes presented on BMDC and MC57G cells. G) The abundance of 45 VACV epitopes on BMDC and MC57G cells. Abundance is presented as the average sum of all timepoints for each epitope. Peptides are coloured by the distance from the log-log regression line to highlight outliers.

Nearly all epitopes were detected within 2.5 hours after infection; however, the range of epitope levels observed on each cell line varied greatly and could differ by more than 1000-fold at a given timepoint (Fig. 1D). At each timepoint, total epitope levels, represented by the sum abundance of all 45 p:MHC-I, were the highest on BMDCs. The most striking difference was the 10-fold lower total epitope presentation on PMFs compared to the other cell types (Fig. 1E).

To identify whether cell phenotype affected how well individual epitopes were presented, we compared the sum abundance of each epitope in each time-course between all four cell types. For individual epitopes presented on phenotypically distinct BMDCs and MC57G cells, we could identify examples where the absolute abundance of an individual epitope had higher, the same, or lower levels on either cell type (Fig. 1F). However, from a multi-epitope perspective, the relative abundance of all 45 epitopes on BMDCs and MC57G cells was significantly correlated (Fig. 1G). Comparing individual epitope levels between all cell types, phenotypically similar dendritic-like BMDC and DC2.4 cells were the most closely related (Supplementary Fig. 1A).

Likewise, epitopes that were more abundant on primary dendritic-like BMDCs compared to primary fibroblast-like PMFs were also more abundant than expected on immortalised dendritic-like DC2.4 compared to fibroblastic MC57G cells (Supplementary Fig. 1B-C). This suggested that similarities in cell type of origin could partially explain how well an individual epitope was presented. A similar comparison between epitopes presented on immortalised cells compared to primary cells was also significant, although not as strong (Supplementary Fig. 1D).

We concluded that VACV p:MHC-I abundance on different cell types *in vitro* was generally correlated, while noting differences in the relative epitope levels between cell types and that big differences can be seen in the case of some epitopes.

### Abundance of mouse and VACV proteins in infected cells *in vitro*

The changes in epitope levels between cell types could be driven by differential mouse antigen presentation machinery expression or altered VACV protein expression. Using a label-free quantification (LFQ) mass spectrometry method, we quantified the abundance of both VACV and mouse proteins from the VACV-infected cells using the lysate of infected cells following p:MHC immunoaffinity purification. LFQ provides a measure of the relative abundance of proteins, without requiring additional protein standards (Cox et al., 2014; Cox and Mann, 2008). 151 VACV proteins were detected across the infected cell types, accounting for approximately 70% of the proteins encoded in the VACV genome. Simultaneously, the abundance of 3588 mouse proteins were detected (Supplementary Table 3).

Principal component analysis (PCA) was initially used to explore protein abundance in this dataset, combining both VACV and mouse proteins (Supplementary Fig. 2), or restricted to either analysis of mouse or virus proteins (Fig. 2A-B). Mouse proteins alone clearly clustered samples by cell type rather than infection time, suggesting changes in the mouse proteome due to infection were minor compared to the differences in protein abundance between cell types (Fig. 2A). Indeed, the stability of the host proteome and the massive abundance of the relatively small number of viral proteins is evident when comparing total amounts of protein with the amounts per-protein between host and viral proteomes (Supplemental Fig 3). In terms of host proteins relevant to epitope abundance, PMFs and MC57G cells were found to have lower abundance of proteins associated with p:MHC-I processing and presentation (Supplementary Fig. 4). In particular, both PMFs and MC57G cells had very low immunoproteasome content and relatively low abundance of proteins that directly form the peptide loading complex in the endoplasmic reticulum (Supplementary Fig. 4D-F). Differences in the relative abundance of antigen presentation machinery were smallest between BMDC and DC2.4 cells (Supplementary Fig. 4C-F).

**Figure 2:**
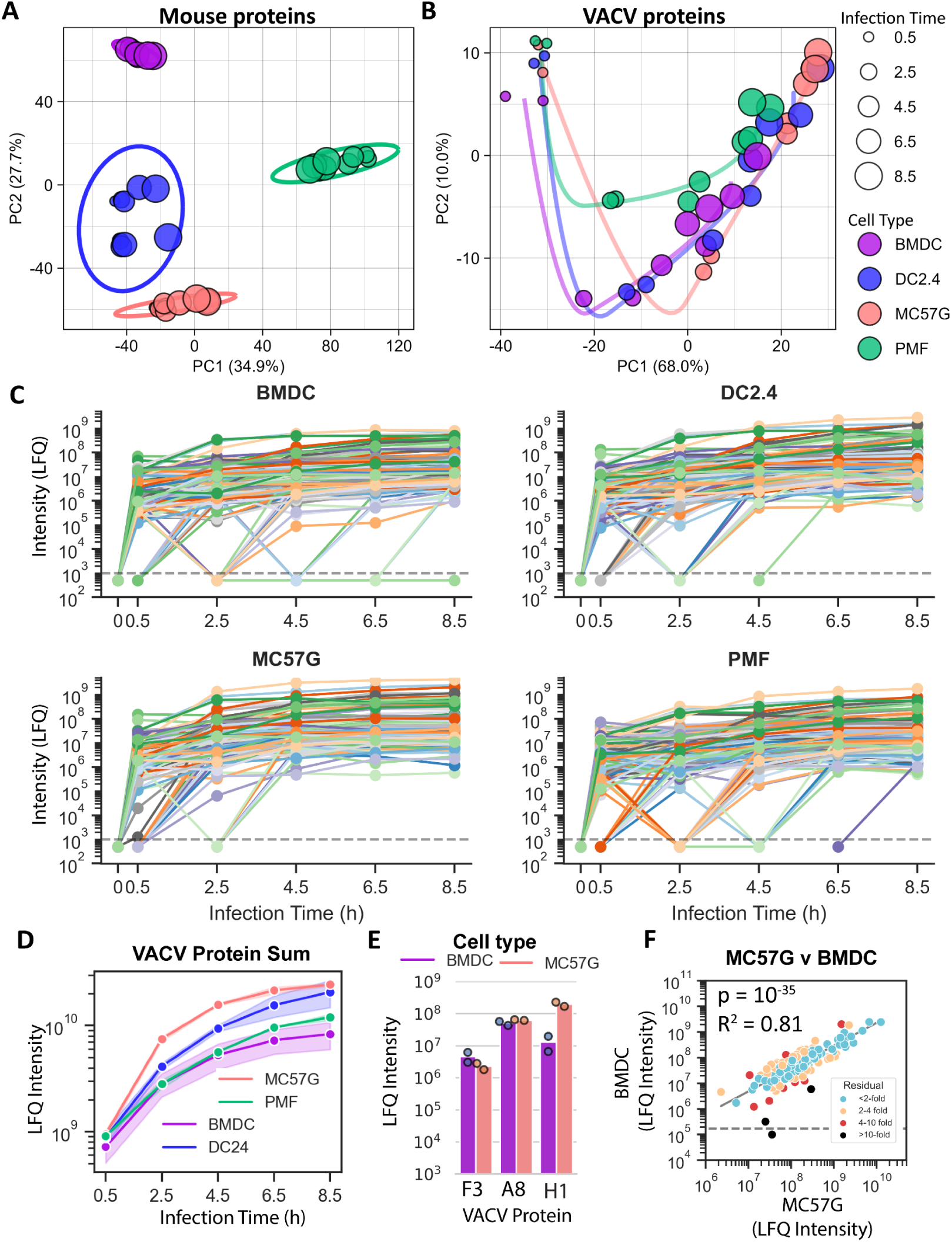
Label-free quantification of mouse and virus proteins *in vitro* on VACV-infected cells over time. BMDC, DC2.4, MC57G and PMF cells were infected with VACV between 0.5h and 8.5hours. Protein abundance was estimated by label-free quantification (LFQ). A,B) PCA analysis of mouse (A) and VACV (B) protein abundance. Clusters identify samples from common cell types using PC1 and PC2 (A) or trace a trajectory of PC1 and PC2 with respect to time projected onto the PC1-PC2 axis (B). C) Average abundance of VACV proteins over time on each cell type at each timepoint. Points below the dotted line represent undetected proteins. Lines are drawn from the time-point before first detection. D LFQ intensities for each cell type at each timepoint. Shaded regions draw the range of values from two independent replicates. E) The sum of protein LFQ intensities over the infection time-course for three VACV proteins. F) Correlation between VACV protein abundance on MC57G and BMDC cells. VACV protein abundance was calculated as the average sum of all timepoints for each protein on log-transformed values. Proteins are colour coded by fold distance from the log-log linear regression line. Points below the horizontal dashed line identify proteins not detected.

In contrast to the mouse proteome, variability in the abundance of VACV proteins was best explained by shifts in protein abundance over time represented by samples following a similar trajectory on the first two principal components regardless of cell type (Fig. 2B). The PCA loadings of each protein suggest PC1 largely represented an increase in the abundance of a VACV protein as the virus replication cycle progressed. Importantly, PC1 and PC2 also separated classes of VACV genes with similar transcriptional kinetics during the virus replication cycle ^16, 51, 52^ (Supplementary Fig. 5).

We next compared the range of individual VACV protein abundance between cell types for each of the 151 detectable VACV proteins over time. VACV proteins at a given timepoint had a wide range of abundances (Fig. 2C). Interestingly, while the abundance of epitopes on PMFs was obviously lower compared to other cell types, total VACV protein abundance on these cells was largely comparable to the other cell types (Fig 2D). Therefore, differences in VACV protein abundance alone could not explain lower epitope presentation on PMFs, but instead inefficient antigen presentation likely reflected the lower abundance of epitope presentation. This is further underlined by the results from infected BMDC, which had the highest levels of epitope presentation, yet the lowest levels of VACV proteins at each timepoint after infection.

For VACV proteins, while some individual proteins were found to have higher or lower abundance across the different cell types (Fig. 2E), overall, the correlation of abundance of VACV proteins between cell types was very tight and notably closer than was seen for epitopes (Fig 2F, Supplementary Fig. 6).

### Most viral epitopes are generated from proteins immediately or soon after translation

To investigate the relationship between epitope presentation and antigen expression kinetics, we compared the absolute abundance of VACV epitopes and the relative abundance of their source proteins at each time point. Previous comparisons of this kind have used a handful of proteins and made visual comparisons comparing the time of peak expression ^24, 40^. Here we were able to examine the presentation kinetics of 41 epitopes. To make a more formal comparison, we calculated the time taken for each to reach the half-maximum levels detected across the time course. Epitopes were classified according to whether epitope half-maximal levels were attained ‘before’, ‘coincident’ or ‘after’ the protein half-maximal levels, defined by whether the epitope half-maximum levels were achieved at least one hour, within one hour, or at least one hour after the protein half-maximum timepoint (Fig. 3A).

**Figure 3:**
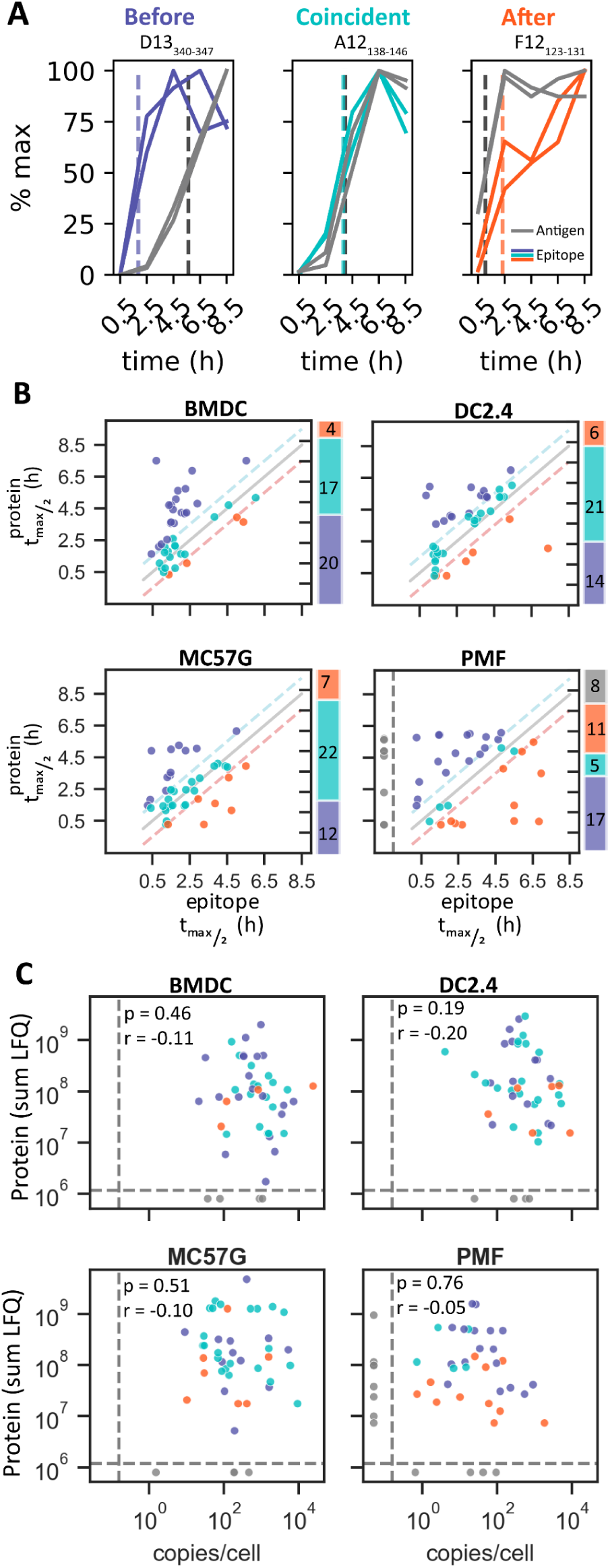
VACV epitope presentation predominantly occurs before or coincident with source protein expression. Epitope and protein abundance was normalized to the maximum value in each replicate over the infection time-course. Four VACV epitopes with undetectable protein levels are excluded from analysis in this figure. A) Representative VACV epitopes on infected BMDCs that show epitope time to half-maximum levels (t_max/2_) occurring before, coincident with, or after the respective protein. Dotted lines identify the epitopes and protein time to t_max/2_. B) Comparison of the time to t_max/2_ for matched epitope and antigen. Diagonal lines identify values representing identical epitope and protein half-life or +/-1 hour, used to classify epitope kinetics as before (blue), coincident (aqua) or after (orange) protein expression or undetected epitope (grey). Bar charts describe the number of peptides classified into each category. C) Comparison of the sum of each epitope and protein abundance (LFQ) over time on VACV-infected cell types. Statistic: Spearman correlation.

Most epitopes reached half-maximal levels either coincident with, or before, the respective source protein. This was particularly evident on infected BMDCs, where nearly all epitopes achieved half-maximal levels within 2.5h of infection and 37 out of 41 epitopes (>90%) were classified as either reaching half-maximal levels before, or coincident with the respective protein antigen (Fig. 3B). Across all cell types, epitopes that were classified into all three kinetic classes could be identified (Supplementary Fig. 7).

The time to reach half-maximal levels for epitopes derived from the same source antigen did appear to be more closely related than for unrelated epitopes (Supplementary Fig. 8), although we could also identify exceptions to this trend. For example, epitopes derived from the same VACV protein A47 were presented either ‘before’ or ‘after’ the detection of the mature antigen (Supplementary Table 1). This shows that the same protein can enter the antigen presentation pathway at more than one point over its lifespan. Likewise, across cell types, the same epitope could be classified into different kinetic classes. These transitions were generally between the ‘before’ and ‘coincident’ kinetic classes, suggesting any cell-type specific changes were mechanistically conservative (Supplementary Fig. 8). However, we also noted a potential polarisation of the epitope kinetics on PMFs into either ‘early’ or ‘late’ classification, resulting in a small increase in the number of late epitopes presented on PMFs (Fig 3B).

### Viral epitope presentation levels cannot be explained by source protein abundance or p:MHC-I affinity

Next, we investigated whether the epitope presentation and respective VACV source antigen levels are related during VACV infection. Across this large set of epitopes and multiple times after infection on four different cell types there was no correlation between protein and epitope abundance (Fig. 3C). These data show definitively that relative amounts of proteins do not predict levels of epitope presentation across a viral proteome. The binding affinity of p:MHC-I has often been thought to be a major determinant of presentation levels. We tested this idea with our data and found that irrespective of cell type, the level of presentation was not significantly correlated with either measured or predicted binding affinity of the peptides for their presenting MHC allomorph (Supplemental Figure 9, Supplementary Table 1).

### Neither epitope nor protein abundance *in vitro* correlates with the size of antigen-specific CD8^+^ T cell responses

Epitope abundance is often suggested to be a primary factor contributing towards the size of peptide-specific CD8^+^ T cell responses during virus infection ^30, 31^. Using the complete set of 45 epitopes, we correlated epitope abundance with the respective peptide-specific CD8^+^ T cell response during VACV infection in C57BL/6 mice following intraperitoneal infection as published previously (Croft et al., 2019). The size of peptide-specific CD8^+^ T cell responses did not correlate with epitope abundance measured on any cell type (Fig. 4A). The absence of a correlation was surprising given the previously described correlation between p:MHC-I affinity and T cell immunogenicity during VACV infection (Croft et al., 2019; Moutaftsi et al., 2006). Having found that epitope abundance did not correlate (see above), for completeness we looked to see if there was a relationship between the relative amounts of viral proteins measured *in vitro* and T cell responses and found that viral protein levels and T cell responses were not correlated (Fig 4B).

**Figure 4:**
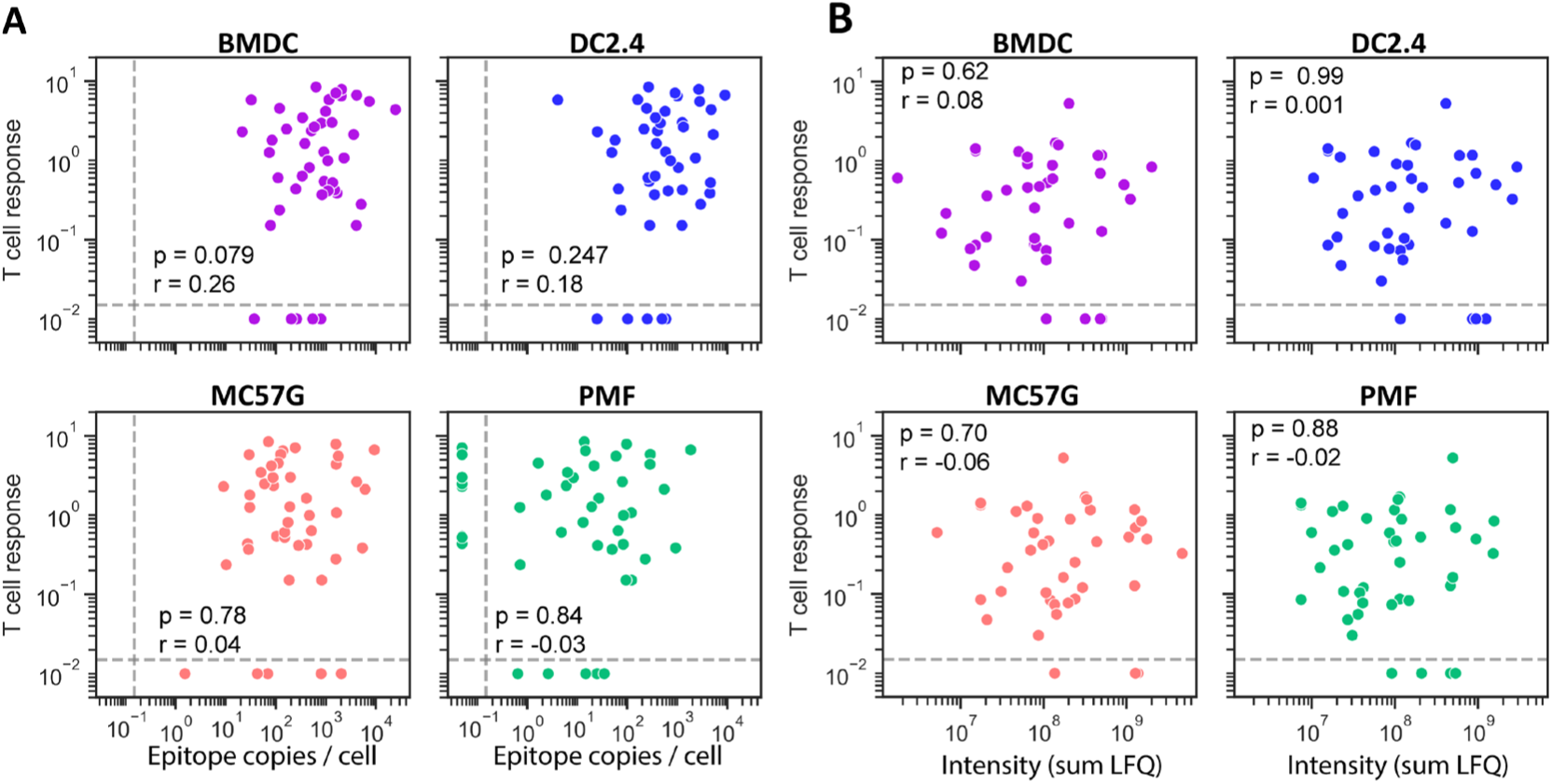
Epitope and protein abundance *in vitro* does not correlate with the size of peptide-specific T cell responses. Epitope (A) and protein (B) abundance was calculated as the average of the sum of values over the time-course from two independent replicates. T cell responses were measured following intraperitoneal infection of C57BL/6 mice (Croft et al., 2019). Statistic: Spearman correlation.

### *In vivo* quantification of epitope abundance and matched CD8^+^ T cell responses

The experiments above were done based on the premise that the actual APC *in vivo* would be displaying p:MHC-I in a similar fashion to at least one of the cell types in our *in vitro* experiments. Looking more closely at the cell-type differences in epitope abundance, we noted an incremental improvement in the correlation coefficient between epitope abundance and T cell responses from fibroblastic cells to those representative of a more likely APC *in vivo*; that is, presentation level on BMDC showed a higher spearman correlation with CD8^+^ T cell magnitude than that on DC2.4, which in turn was higher than seen for MC57G and PMF. For this reason, a new strategy was devised in which we set out to measure VACV epitope abundance *in vivo* in an experimental infection model that we could match for the measurement of CD8^+^ T cell responses. Specifically, we a) measured VACV epitopes on the spleens of mice 24 hours after an intravenous injection with VACV and b) reassessed the CD8^+^ T cell responses to the set of epitopes after an intravenous infection because immunodominance hierarchies can be altered by route of infection ^46^ (Fig. 5A).

**Figure 5:**
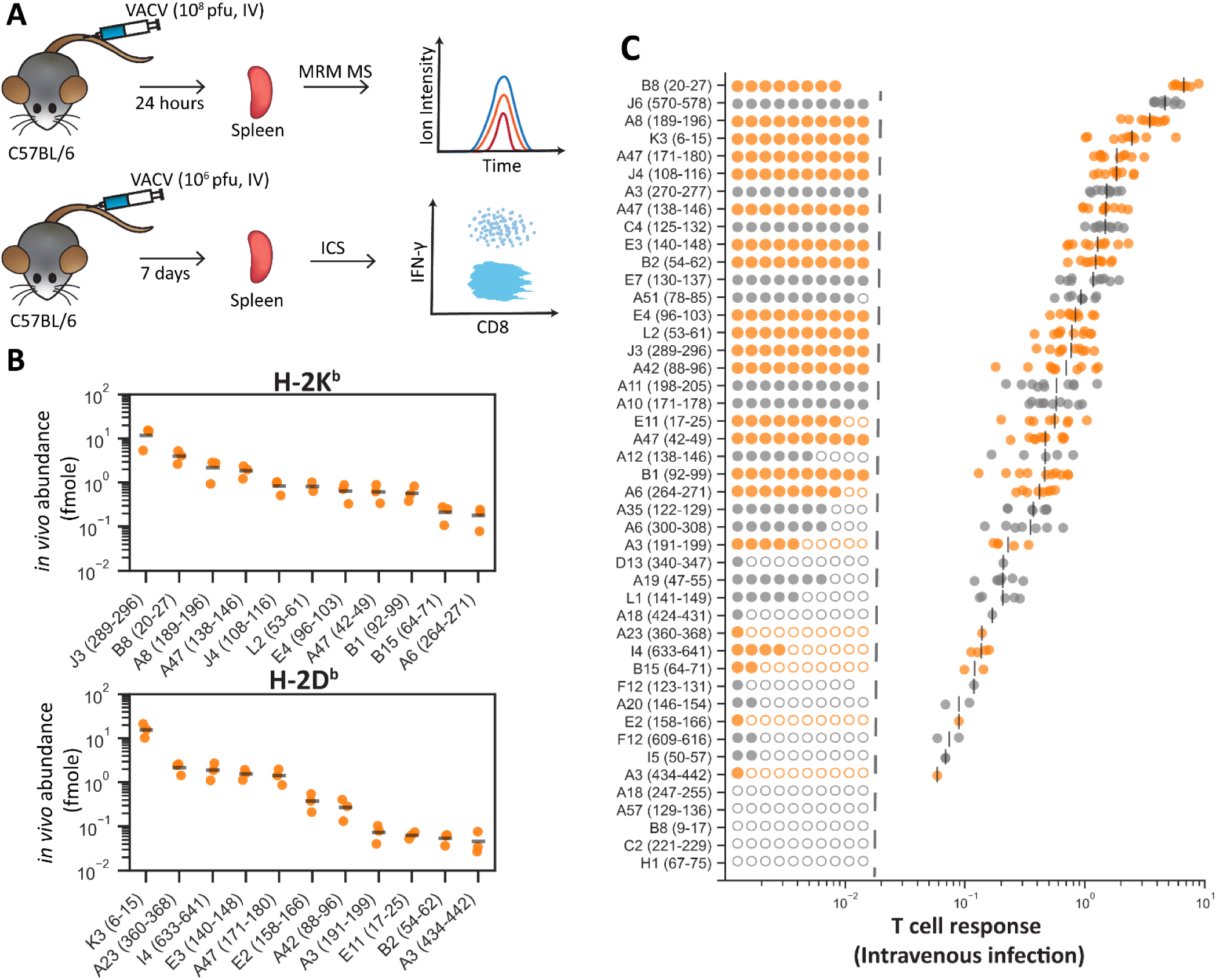
A shared experimental model for *in vivo* quantification of epitope abundance and measurement of peptide-specific CD8^+^ T cell responses. A) C57BL/6 mice were infected with 10^8^ pfu VACV by intravenous infection. 24hours p.i. VACV epitope abundance from the spleen was measured by MRM MS. B) C57BL/6 mice were infected with 10^6^ pfu VACV by intravenous infection. 7 days p.i., the size of peptide-specific CD8^+^ T cell responses to 45 VACV-specific epitopes were measured by intracellular cytokine staining. C) Abundance of peptide presented on H-2K^b^ and H-2D^b^ from the spleen of three infected mice. Peptides not detected in any mouse are not shown. D) Peptide-specific CD8^+^ T cell responses measured by ICS following intravenous infection. The set of epitopes detected by MRM MS in the spleen (C) are coloured in yellow. Filled and empty circles left of the dashed line identify the number of mice a peptide-specific T cell response was immunogenic (filled) or non-immunogenic (empty). Where a response was determined to be immunogenic, the size of the response is shown on the right. Bars represent the mean immunogenic T cell response.

Using MRM MS to measure the abundance of each epitope, 22 out of the 45 epitopes investigated in this study were detected in the spleen after infection, spanning over a 300-fold range in abundance (Fig. 5B). Although approximately 50% of epitopes we investigated were not detected *in vivo*, many of these peptides generate CD8^+^ T cell responses and it is likely that their absence highlights the technical challenges of detecting very low quantities of virus-derived p:MHC-I rather than the failure of these peptides to be presented *in vivo*.

Next, the magnitude of peptide-specific CD8^+^ T cell responses in the spleen seven days after intravenous infection were measured by restimulation of cells with peptide followed by staining for intracellular IFNγ (Fig. 5D). Consistent with previous virus infection models, some peptide-specific CD8^+^ T cell responses were detected only in a fraction of mice (Croft et al., 2019; Wu et al., 2019). In these cases, the average peptide-specific responses were calculated only from mice where an immunogenic peptide-specific response was detected (Croft et al., 2019). These data also highlight the diversity of the CD8^+^ T cell response, in spite of marked immunodominance, with responses to around six peptides contributing half of the total measured anti-viral response and >20 peptides accounting for over 90% of the response (Supplementary Fig 10).

### Epitope levels measured *in vivo* correlates with the CD8^+^ T cell immunodominance hierarchy

Using the set of 22 epitopes detected *in vivo*, we examined the relationship between epitope abundance and the size of peptide-specific CD8^+^ T cell responses following intravenous challenge. All peptides in this dataset were immunogenic and therefore there was no observable threshold in epitope abundance below which CD8^+^ T cell activation would not occur.

For the first time, we identified a significant correlation between epitope abundance *in vivo* and peptide-specific CD8^+^ T cell responses during VACV infection (Fig. 6A). Mirroring the relationship observed on infected cells *in vitro,* p:MHC-I affinity, whether measured, or predicted was not statistically correlated with epitope abundance (Fig. 6B).

**Figure 6:**
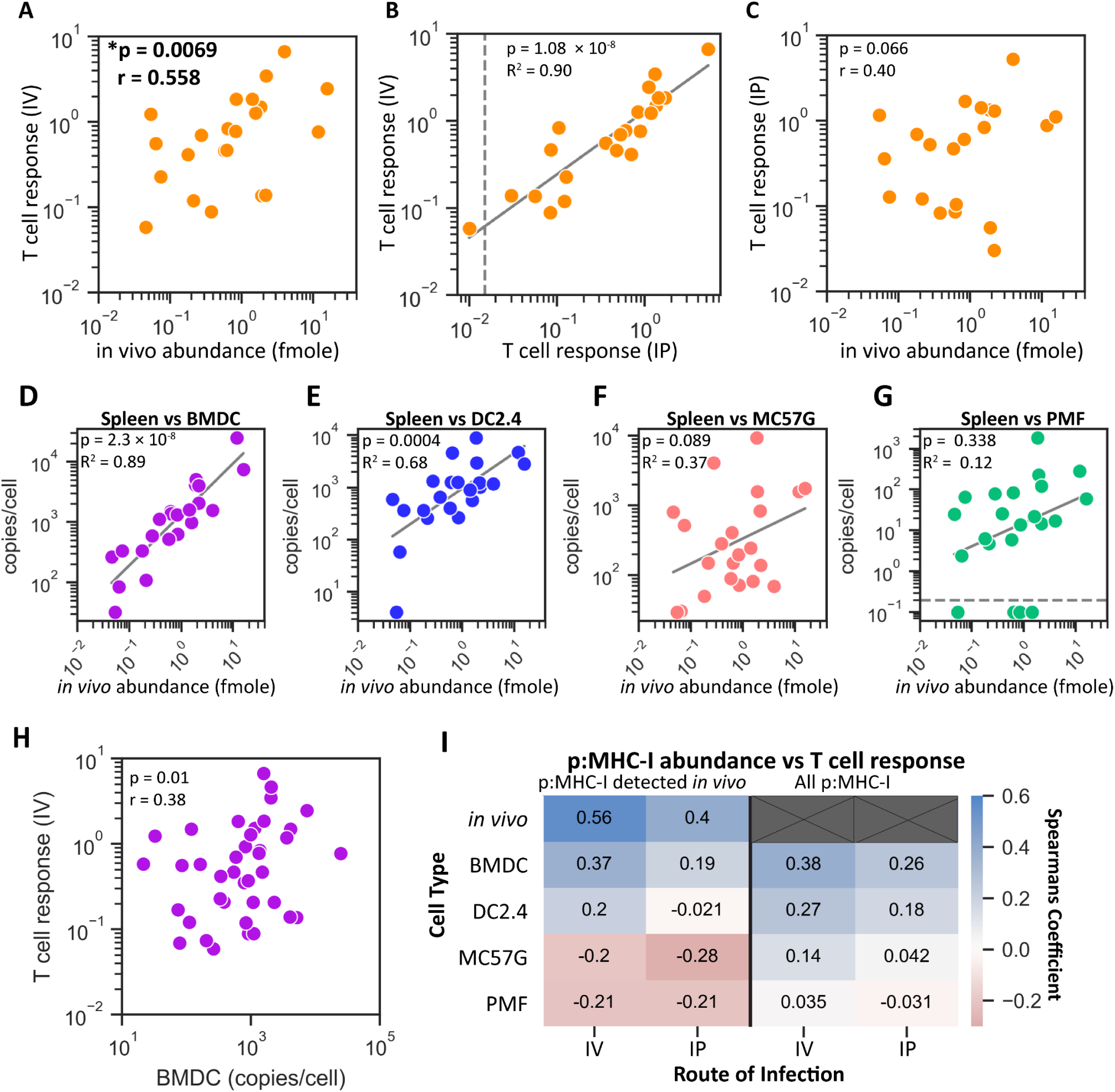
*In vivo* measurement of epitope abundance correlates with peptide-specific CD8^+^ T cell responses following intravenous infection. A) Correlation between the average epitope abundance *in vivo* and the average immunogenic peptide-specific CD8^+^ T cell responses following intravenous infection with VACV. B) Comparison of peptide-specific CD8^+^ T cell responses in C57BL/6 mice following intraperitoneal T cell responses (Croft et al., 2019) and intravenous T cell responses measured in this study. C) Comparison of peptide-specific CD8^+^ T cell responses following intraperitoneal T cell responses in C57BL/6 mice and epitope abundance measured *in vivo* (Croft et al., 2019, current publication). D-G) Correlation between epitope levels measured *in vivo* and on VACV-infected cells *in vitro*. H) Comparison of epitope abundance measured on infected BMDC cells *in vitro* and peptide-specific CD8^+^ T cell responses following intravenous infection. I) Degree of correlation between epitope abundance and peptide-specific CD8^+^ T cell responses. Comparisons include the set of 22 epitopes detected *in vivo,* or the complete set of 45 epitopes. Statistics: Spearman correlation: A,B,H,I; Pearson correlation: C-G

To understand why we identified a statistical correlation between epitope abundance *in vivo,* but not *in vitro*, we investigated whether this might be due to a shift in the magnitude of T cell responses when examined after an intravenous infection, differences in epitope abundance *in vitro* and *in vivo*, or a combination of both.

The CD8^+^ T cell immunodominance hierarchies determined via intraperitoneal (Croft et al., 2019) and intravenous infection were very closely correlated (R^2^=0.90), consistent with only minor shifts in the CD8^+^ T cell immunodominance hierarchy from the altered route of infection (Fig. 6B). The most notable difference was the reduced dominance of the immunodominant B8_20_ epitope relative to other epitopes, recapitulating published observations (Lin et al., 2013). This difference in magnitude of the T cell responses between infection routes alone was enough to reduce the significance of the correlation with the level of epitope presentation measured in vivo, reducing the quality of the relationship than then when the T cell responses were measured after intravenous infection (Fig. 6C).

Epitope abundance *in vivo* and on cells *in vitro* was also strongly correlated, most notably for BMDC cells *in vitro* but progressively less so for DC2.4, MC57G and was weakest for PMF in a similar manner to our observations when exploring presentation levels and T cell responses (Fig. 6D-G). This stepwise erosion of the relationship between epitope levels measured *in vivo* and *in vitro* is consistent with the previous observation that dendritic cells and macrophages are the primary cell types infected with VACV in the spleen ^53^. We concluded that epitope abundance measured *in vitro* is broadly representative of p:MHC-I abundance *in vivo,* although this is most true when the antigen presenting cell type *in vitro* reflects the biologically relevant cell type presenting p:MHC-I *in vivo.* The relationship between presentation on BMDC and in the spleen *in vivo* was so close that we re-examined whether the epitope amounts presented on these cells might correlate with the T cell response, this time using the data generated by intravenous infection. When we restricted our analysis to the 22 epitopes identified *in vivo,* the correlation was not statistically significant. However, when we expanded the data to include all 45 epitopes this relationship now achieved the p<0.05 level required for statistical significance (Fig. 6I). This result encouraged us to go back to the epitope levels quantified *in vitro* to ask whether measuring these at any particular time after infection might be best to predict immunogenicity (Supplementary Fig 11). We found that epitope levels measured at 2.5 hours after infection, the second time point in the experiment, were the best correlate of immunogenicity on DC cell types, being superior to the overall presented amounts determined by summing presentation across all times measured.

Together these observations demonstrate that levels of epitope presentation are a general correlate of the size of T cell responses, however the biology of CD8^+^ T cell priming is complex and this relationship can be obscured by other variables. As a result, the significance of the correlation only emerges if confounding variables are reduced by very close matching the models used to measure epitope abundance and the T cell response and/or a very large number of epitopes is examined.

## Discussion

Almost twenty years ago, the indiscriminate use of poorly-defined epitopes to characterise CD8^+^ T cell responses was highlighted in a review article as a ‘dirty little secret’ in the field of antigen presentation and attempts to relate presentation with immunogenicity ^54^. This review proposed a system of escalating criteria by which epitopes might be ranked for how well they have been defined. Key to achieving a higher ranking was the use of mass spectrometry to identify the relevant peptide being presented on infected cells, with the highest rating reserved for the same being achieved directly *ex vivo*. Advances in mass spectrometry to detect presented epitopes on cells *in vitro* has become more common, but attempts at quantification remain rare ^24^. Even more scarce has been detection of peptides directly from tissue of infected animals but prior to this study, systematic quantification had not been achieved ^55, 56^. Further, comprehensive studies that combine *in vitro* and *in vivo* quantification of epitope presentation across a large set of epitopes, with the aim of finding general correlates of immune responsiveness have been lacking and this is the gap we fill with our data and analysis.

In this study, we set out to address four fundamental questions: 1) the extent to which presentation of epitopes differs across antigen presenting cell types; 2) whether antigen levels predict epitope levels; 3) the kinetic relationship between viral protein production and epitope presentation and finally 4) whether epitope presentation levels are a significant driver of immunogenicity. First, we found that when considered as varying over orders of magnitude, epitope presentation levels broadly correlated between all infected cell types *in vitro* (Fig 1; Supplementary Fig. 1). This was most striking for PMFs, in which epitope abundance correlated with abundance from all other cell types despite the globally lower presentation levels of all epitopes. Having noted this overall result, individual epitopes could be found that differed in abundance by more than 10-fold between different cell types. This helps to explain disparate findings in the literature where fewer peptides were examined and is reminiscent of data published for influenza A virus, though the correlation shown here for VACV epitopes was stronger ^24^. The differences we do observe are most likely related to antigen processing rather than in virus infection, because the correlation of viral protein levels between pairs of cells was higher than for epitopes (Fig 2). Again, PMFs provide a clear example of this effect given the similar levels of overall viral protein abundance compared with the other cell types, but lowest levels of proteins associated with antigen processing and presentation and poor levels of epitope presentation (Supplementary Fig 6). The more subtle shifts in the relative abundance of epitopes, such as between BMDC and MC57G cells, could arise from the altered biases in the processing of antigen. This includes altered proteolytic biases of immunoproteasomes (in BMDCs) compared to constitutive proteasomes (in MC57G), which may preferentially increase the processing efficiency of particular epitopes ^12, 13^. The presence or absence of immunoproteasome subunits has been suggested to result in major changes to the immunopeptidome, including the generation or destruction of epitopes during antigen processing ^12, 57, 58^. However, a systematic, quantitative analysis by Mishto, Liepe ^14^ measuring polypeptide degradation *in vitro* by different proteasomal isoforms indicated that the functional differences between constitutive and immunoproteasomes are quantitative rather than qualitative ^14^. The latter observation better aligns with the shifts in epitope abundance between cell types observed in this study, although the extent to which this can be attributed to the level of immunoproteasome activity is unknown. Finally, it should be noted that for viruses that have very different replication kinetics, including abortive infections in some cell types, it is possible that epitope presentation may differ much more across cell types than we find here. This highlights the need to understand not only antigen presentation, but also the unique biology of any virus being studied, although our data suggests that where infection is comparable, presentation will be similar across epitopes.

Second, as noted above, the kinetics and abundance of individual proteins in the viral proteome had differing characteristics than their representation within the immunopeptidome. Therefore, it was unsurprising that in our experiments source protein amounts failed to correlate with the abundance of epitopes. This was found for all timepoints after infection examined and for all cell types. It also confirms similar experiments using influenza virus but contradicts some suggestions based on presentation of self-antigens that suggest protein abundance is an important variable ^24, 59–62^. This might represent a difference between viral and self-antigens, but may also reflect different methodologies: we are directly measuring epitope abundance where other studies infer abundance from the likelihood of identification or RNAseq data ^59–61^. Self-antigen studies also have much larger data sets allowing even weak associations to be identified. Given the large set of epitopes and proteins we have available here for VACV, it seems unlikely that protein abundance is going to be found to be helpful in predicting epitope presentation levels for any other virus.

Third, as suggested in studies of influenza virus and Epstein-Barr virus, we find here with our set of more than 40 epitopes that most VACV epitopes are processed from their source protein as quickly as these proteins are translated ^24, 63^. Adding some nuance to this finding is the change in extent to which this occurs when looking at the data from professional APCs and particularly the BMDC, compared with the fibroblastoid cells, with the primary mouse fibroblasts being most different. In the BMDCs, less than a tenth of the source proteins reached their half maximal level before the epitope, but for PMFs this is a third. It is tempting to speculate that the specialisation of DCs for presentation goes beyond the immunoproteasome and superior levels of protein components of the antigen processing machinery and extends to an ability to sample newly translated proteins more effectively. Why some epitopes are not rapidly presented coincident with translation of the source antigen is not clear, although this may be associated with the rate of protein translation, or turnover as has been suggested for self-peptides ^60^. In favour of the idea that it is a property of the viral protein, where we have epitopes derived from the same source antigen they tend to have a similar epitope:antigen kinetic profile and this is maintained across cell types. A limitation of our analysis and data is that we measure only ‘bona fide’ proteins, which will miss any DRiPs or other short-lived or non-canonical products of translation that may be sources of epitopes.

Finally, we have assessed the extent to which the abundance of epitope presentation is correlated with their immunogenicity, across a wide range of specificities. This is another area where previous studies are split between finding correlations or not, across a range of viruses and number of epitopes under investigation ^24, 32, 33, 35, 40^. The contribution that we make here is two-fold: 1) we answer this question with a large number of epitopes such that our data are likely to be robust and generalizable and 2) by making multiple measurements of epitope abundance *in vitro* and *in vivo*, it is clear why the literature has opposing answers to what should be a fundamental question.

The magnitude of epitope-specific T cell responses is subject to multiple factors unrelated to antigen presentation including the number of naïve T cell precursors, TCR affinity for an epitope and susceptibility to immunodomination ^28, 32, 64–67^. These complex multi-factorial contributions towards T cell responses add significant noise to attempts to correlate factors associated with antigen presentation and the magnitude of responses. Further, while epitope affinity for MHC-I has been viewed intuitively as a factor that underlies the abundance of epitope presentation, our data would suggest that this is not the case: affinity and abundance were not closely correlated on cells infected *in vitro* or the amount of epitope presented in the spleen. These suggest that affinity and abundance have separate, if overlapping contributions to immunogenicity as has been suggested previously ^24^. The most likely role played by affinity beyond ensuring adequate presentation on the surface of cells is in stabilising p-MHC-I:TCR interactions. In the case of influenza and lymphocytic choriomeningitis viruses, IC_50_ of peptide for MHC-I was inversely correlated with T cell responses and a similar result was seen over a larger range of epitopes than have been studied here for VACV ^4, 24, 28^. We were not able to see this correlation here, however we chose a subset of epitopes with a relatively high affinity for MHC-I (geometric mean = 2.5 nM) from the total number of >150 VACV epitopes (geometric mean = 18nM). This might mean that that IC_50_ is less important above an affinity threshold, but none-the-less affinity of peptide for MHC-I seems likely to be a factor that influences both antigen presentation and T cell activation. Given these other sources of variability from the T cell side, it is not surprising that a large number of epitopes is required for a correlation between abundance and T cell responses to be found.

The variability in the literature is likely to derive from studies with too few epitopes, but also with the differences in how epitope abundance was measured, including cell and infection type and how closely this mimics presentation *in vivo* on the DCs that prime the CD8^+^ T cell response. The extent to which this matters will be dependent on the biology of the virus. For VACV, we show a sequence of apparently subtle shifts in presentation amounts looking across our data from *in vivo* to infected fibroblasts. Had these studies been published separately, we would have confidently interpreted our results in a study of fibroblasts as finding no correlation between abundance of presentation and T cell responses and then the opposite in a second study in which we measured abundance *in vivo*. Presenting these together makes it clear that it matters how epitope abundance is measured, with *in vivo* being by far the best.

The extent to which the infection model matters for understanding antigen presentation will depend on the virus or other pathogen used. In the case of our study, we can see the biology of VACV infection and immunity reflected in the data. For example, confocal microscopy of the spleen from infected mice indicate that macrophages and dendritic cells are the primary cell types infected in the spleen during VACV infection ^68^, and confirm infected bone-marrow-derived dendritic cells are the predominant cell type that prime CD8^+^ T cells during VACV infection ^69^. This is mirrored in the close correlation we observed between epitope levels from infected spleen and on dendritic-like cells infected *in vitro*, and the loss of significant correlation when comparing with fibroblasts. Similarly, there is substantial evidence that infection of DCs and direct-presentation of epitopes during VACV infection is the dominant mode of priming for most CD8^+^ T cells ^27, 31, 47, 70^. This too is in alignment with abundance of presentation in spleen and infected DCs being very similar. It is instructive to compare results here with that for influenza virus, which uses both direct and cross priming, with one particularly dominant response (PA_224_) being largely cross primed ^71^. Interestingly when antigen abundance on infected cells and on DCs cross-presenting influenza proteins were correlated with CD8^+^ T cell responses, cross presented amounts were the best correlate unless responses to PA_224_ were discarded ^24^. This illustrates again that the biology of the virus matters in assessments of epitope abundance when used to understand immunogenicity.

Overall, our study highlights that *in vivo* antigen presentation can be recapitulated to a large extent *in vitro*. However, significant correlations between CD8^+^ T cell immunogenicity and epitope abundance were most clear measuring epitope levels directly from cell types *in vivo* expected to directly prime CD8^+^ T cells during an infection. Going forward, similar *in vivo* models should be considered the gold-standard approach to assess epitope abundance. Further, the quantification of epitopes from the splenocytes of infected mice represents a significant technical advance and sets a new benchmark for viral immunopeptidomics. Clearly, much has been said about antigen load and epitope presentation in the context of virus infections based entirely on indirect measures. Our quantitative approach to measure epitope abundance from the spleen of infected mice is the first to directly identify a correlation between epitope levels *in vivo* and the magnitude of peptide-specific T cell responses. Finally, our systematic approach that used a large set of epitopes spanning orders of magnitude in their levels of presentation and immunogenicity gives us confidence that the results here represent general correlates that will be broadly relevant across pathogens.

## Methods

### Mice

All mice were obtained, housed and bred from the Australian Phenomics Facility (Canberra, Australia). Mouse were housed and all experiments were conducted according to the ethical requirements under approvals A206/45, A2020/01 and A2023/09 from the Australian National University Animal Ethics and Experimental Committee. Specific pathogen-free Female C57BL/6 mice over 8 weeks of age were used for all experiments except where indicated.

### Cell lines

All immortalised cell lines, including DC2.4 and MC57G cells were cultured in T-175 or T-75 flasks containing in DMEM + 10% FCS + L-glutamine and passaged every 3-4 days using trypsin to detach adherent cells. Cells were grown in incubators at 37 °C in 5% CO_2_.

### Culturing primary Bone-Marrow derived Dendritic cells

Bone-marrow derived dendritic cells (BMDCs) were prepared and cultured over 6 days from female mice bone-marrow cells as described by Madaan, Verma ^72^. C57BL/6 mice were culled by CO_2_ asphyxiation. The femur was isolated, and further cleaned with 70% ethanol. The femur was cut above the knee-joint and below the epiphysis and the contents of the femur were flushed with 2mL PBS using a 1mL insulin syringe (25G × ½ needle) and collected in a 50mL centrifuge tube.

Bone-marrow cells were suspended in 20mL cold PBS and centrifuged for 8 min (250 × g). Supernatant removed, and cells washed twice with PBS and resuspended in BMDC media (DMEM; high glucose, 2mM L-glutamine, 10% FCS, 1% Penicillin/Streptomycin, 55µM 2-mercaptoethanol, 50 ng/mL rGM-CSF). 8mL of cells containing 10^7^ cells were added to a 90-mm petri dish (non-tissue culture grade). 4mL of BMDC media was replaced on days 2,4 and 6. Petri dishes were placed in an incubator at 37 °C, 5% CO_2_ and 95% humidity for 3 days. On day 8, LPS was added to culture dishes to a final concentration of 1ug/ml, incubated for two hours and then cells were harvested for infection with VACV. GM-CSF (50 ng/ml) was maintained in all media, including throughout virus infection.

### Culturing Primary Murine Fibroblasts

To generate primary mouse fibroblasts, 3-5 day old male and female neonatal mice were culled by decapitation, tails removed and bodies skinned. Tails and skin were placed in a 90mm petri dish with 4mL warm PBS. The skin was further finely cut with a scalpel blade (Swann-Morton) and split into two 15mL Falcon tubes. 5mL of Collagenase IV (Worthington) was added per tube and incubated for 25 minutes at 37°C. Samples were centrifuged for 5 minutes at room temperature, 205×g and supernatant removed. Cells were washed in 5mL PBS and centrifuged again before the supernatant removed. Cells were incubated in 5mL of 0.05% trypsin for 20 minutes at 37 °C and centrifuged again. The cell pellet was resuspended in 5mL of fibroblast media (DMEM, 10% FCS, 1% (v/v) MEM NEAA, 1% (v/v) penicillin streptomycin, 1% (v/v) L-glutamine) and transferred to a single Nuclon Delta-coated, filter-capped 175 cm^2^ tissue culture flask (Thermo-Fisher Scientific). 20mL of additional fibroblast media was added to each flask and incubated at 37°C in 5% CO_2_. For the first four consecutive days, media was carefully replaced.

On day four, cells were washed in PBS and treated with 0.05% trypsin for 3-5 minutes. Trypsin was inactivated with 10mL fibroblast media and cells separated into four 175cm^2^ tissue culture flasks with an additional 20mL fibroblast media. Once cells reached confluency, cells were washed, trypsinized and seeded into eight 175cm^2^ flasks. After this step, cells were grown to confluency and split 1 in 2 over two passages to generate 32 T175 flasks.

### Generation of VACV virus

VACV strain WR (equivalent to ATCC VR-1354 = WR (NIH TC-adapted); GenBank: AY243312.1) was sourced directly from Dr B Moss (NIH) and is referred to throughout as VACV. VACV was grown in BSC1 cells grown to confluency in D10 media. Each flask was inoculated with 3 × 10^6^ pfu VACV in D2 media. Flasks were incubated for 3 days at 37°C. Cells were harvested, and lysed using a glass homogenizer (Wheaton) and then rapidly frozen and thawed 3 times. VACV stocks were purified by sucrose cushion purification using a 36% sucrose solution.

### Titration of VACV virus

VACV *WR* (referred to as VACV throughout) was diluted in 10-fold serial dilutions before adding to 6-well plates with a BSC-1 cell monolayer. After 90 min incubation at 37^p^C with 5% CO_2_, media was replaced with 2mL per well of 0.4% Sodium carboxymethyl cellulose (CMC) and 2% FCS. Plates were incubated at 37°C for 3 days. Virus plaques were counted by staining the cell monolayer with a crystal violet staining solution to determine virus titre.

### Infection of Mice

C57BL/6 mice were infected with VACV via an intraperitoneal injection or intravenous injection. Mice were infected with 10^6^ pfu for experiments directly measuring T cell responses, and 10^8^ p.f.u to directly measure VACV p:MHC-I presentation *in vivo.* Mice were monitored daily after infection.

### Infection of cell lines

BMDC, DC2.4, primary murine fibroblast (PMF) or MC57G cells were grown or cultured as described. Adherent cells were detached with trypsin, and resuspended in D10 media. Cell viability and concentration was measured using trypan blue staining, and 1 × 10^8^ cells were prepared for each time-point. Cells were washed in 20mL D0 media and resuspended in 1mL in a 14 ml round bottom tube (BD). 1mL of virus was added to the cells at an MOI of 5, and the tubes incubated in a 37°C shaking incubator (250rpm for 30 minutes). After the initial incubation, cells were transferred to 40mL of D2 in a 50mL centrifuge tube and incubated at 37°C for the respective time (0, 2, 4, 6 or 8 hours). The tubes were gently rolled with a MACSmix™ Tube rotator. At each timepoint, the sample was centrifuged for 5 minutes (1500rpm, 4°C) and washed with cold PBS. Next, virus was inactivated in 1mL of PBS containing 1 μg/mL trioxsalen (Sigma-Aldrich), followed by ultraviolet (UV) irradiation (Vilber Lourmat VL-215.L 365nM UV) for 20 minutes on mice. Every 5 minutes, the samples were mixed gently. After UV irradiation, each sample was transferred to 50mL PBS and centrifuged for 5 minutes. PBS was aspirated and samples rapidly frozen using a dry ice ethanol bath. Samples were stored at –80°C until further use.

### Intracellular Cytokine Staining

Female C57BL/6 mice were infected with 10^6^ pfu of the specified VACV strain by intraperitoneal or intravenous injection. Seven days p.i, mice were culled and the spleen isolated for analysis as previously described by Flesch, Hollett ^73^. Briefly, 1 × 10^6^ splenocytes were plated into round-bottom 96-well plates and incubated with a final concentration of 10^-7^ M synthetic peptide (Supplementary Table 2) at 37°C with 5% CO_2_. After 1 hour incubation, 50 µg/mL brefeldin A was added to a final concentration of 5 µg/mL and plates were incubated for a further 3 hours. After incubation, the splenocytes were spun at 4°C to remove the medium and stained with an α-CD8 (PE) antibody in a total volume of 50 μL per well, washed twice with FBS-PBS and then fixed with 1% paraformaldehyde (PFA). Cells were centrifuged and media removed. α-IFN-γ (APC) antibody was diluted in FBS-PBS containing 0.25% (w/v) saponin. 50 μL was added to each well and plates were incubated overnight at 4°C. Finally, cells were washed in FBS-PBS three times and analysed by flow cytometry.

To determine the size of a peptide-specific CD8+ T cell responses, or the proportion of effector CD8+ T cells that were immunogenic, cells were analysed using a BD LSRII flow cytometer. For each sample, at least 50,000 single, CD8^+^ T cells were identified by gating for live singlet lymphocytes on forward scatter (FSC-A) × side scatter (SSC-A), followed by a FSC-A × FSC-H and SSC-H × SSC-W singlet gating strategy and CD8^+^ × IFNγ to identify the proportion of IFNγ^+^ CD8^+^ T cell response. Background responses were determined for samples incubated with mock DMSO treatment without peptide, and were subtracted from the values presented.

The average peptide-specific CD8+ T cell response was calculated by considering only mice where the peptide was determined to be immunogenic, defined by a response at least 3-times above the standard deviation of the background CD8^+^ T cell responses.

### Solubilising lyophilized peptide stocks

Isotopically-labelled AQUA peptides were solubilised in 100% DMSO to a final concentration of 5mM. The concentration of peptide was measured using a Direct Detect® spectrophotometer (EMD Millipore) as per manufacturers instructions. Peptides were diluted to 5 pmoles/μL in MS Buffer A.

### Cross-linking immunoaffinity columns

To prepare immunoaffinity columns for MHC-I purification, 10mL of 10% (v/v) acetic acid was incubated in poly-prep chromatography columns (Bio-Rad) for 20 minutes. Each column was washed with 10% (v/v) acetic acid and rinsed with Milli-Q water before drying. Protein A sepharose was added as 50% slurry to 20% ethanol, mixed and transferred to a column. Each column was rinsed with 10 column volumes of PBS. Antibody (Y-3, specific for H-2K^b^; 28-14-8S, specific for H-2D^b^) to capture MHC-I complexes was diluted in PBS, resin at a concentration of 10 mg per 1mL of resin and then incubated for 1 hour at 4°C while gently rotating, before transferred back to the column. 10 column volumes of borate buffer (50 mM Boric acid, 5 0mM KCl, 3.97mM NaOH) was used to wash the unbound antibody, followed by 10 column volumes of 0.2 M Triethanolamine, pH 8.0. Bound antibody was cross-linked to protein A sepharose using 40mM DMP cross-linker for 1 hour at RT. Crosslinking was stopped by the addition of cold 0.2 M Tris, pH 8. Each column was rinsed with 10 column volumes 0.1 M citrate buffer (0.1M Citrate, pH 3.0), followed by 10 column volumes of borate buffer. Prepared columns were stored at 4°C before use.

### Homogenisation of infected cells

To homogenise infected cells and spleens for p:MHC-I immunopurification, spleens were snap frozen, ground in a Retsch Mixer Mill MM 400 under cryogenic conditions and resuspended in a solution to deactivate and solubilise p:MHC-I using a lysis buffer containing 0.5% IGEPAL CA-630 (Sigma), 50nM Tris-HCl (pH 8.0), 150mM NaCl, and a Complete protease inhibitor tablet (Roche Molecular Biochemicals).

### Immunoaffinity purification of p:MHC-I complexes

Purification of p:MHC-I columns were modelled on the protocol described by Purcell, Ramarathinam ^74^. 1.5mg of cross-linked antibody and protein A sepharose slurry was added to a clear polyprep column. Each column was washed with 10 column volumes of Wash buffer 1. Cell lysate was added to a pre-column, containing protein A sepharose that did not contain cross-linked antibody, that removed contaminants binding non-specifically to protein A. Next, flowthrough from the pre-column was dripped directly onto an immunoaffinity column containing α-H-2K^b^ antibody bound to protein A sepharose. To ensure efficient binding of p:MHC-I to the antibody, the lysate was mixed and incubated for 5 minutes at 4°C. The lysate passed through the column by gravity flow and added to the same column to pass the lysate again. Flow-through was added directly onto an α-H-2D^b^ column, mixed with the cross-linked protein A sepharose slurry, and passed through the column as described for H-2K^b^. Each immunoaffinity column was washed to remove unbound material and contaminants sequentially with Buffer 1 (0.005% v/v IGEPAL, 50 mM Tris, 150 mM NaCl, 5mM EDTA, 100 µM PMSF, 1µg/mL Pepstatin), Buffer 2 (50 mM Tris, 150 mM NaCl), Buffer 3 (50 mM Tris, 450 mM NaCl) and Buffer 4 (50 mM Tris). Antibody-capture p:MHC-I was eluted with 5 column volumes of 10% (v/v) acetic acid from each column separately. Finally, 50fmoles of AQUA peptide were added to the eluate.

### HPLC purification and fractionation

Eluate from immunoaffinity purification contained peptides, the MHC-I heavy chain and β2-microglobulin. To separate these molecules, the eluate was fractionated as described by Croft, Smith ^40^ using reverse-phase HPLC on a 4.6 mm internal diameter× 50 mm long reversed-phase C18 HPLC column (Chromolith Speed Rod, Merck) using an AKTAmicro HPLC system (GE Healthcare). HPLC Buffer A (0.1% TFA, 99% Optima MS-grade water) and HPLC Buffer B (0.1% TFA, 80% Acetonitrile, 19.9% Optima water) were used in the phase buffer system with a flow rate of 1 mL/min. To separate peptides from the MHC-I heavy chain, and β2m, and fractionate the peptides by hydrophobicity, peptides were separated across a gradient of HPLC buffer B from 2% to 45% over 20 minutes. 500μL of each peptide fraction was collected in LoBind Eppendorf tubes and concentrated by vacuum evaporation. Peptide fractions were pooled using a concatenation strategy, maximising retention time distribution across the subsequent RP-HPLC used during mass spectrometry detection.

### Preparation of cell lysate for proteomics analysis

Flow-through from each immunoprecipitation experiment was kept for proteomics analysis. ∼50 µg of lysate material was treated with the reducing agent TCEP at a final concentration of 5 mM and heated to 60 °C for 30 minutes before being loaded onto a FASP protein digestion column (Abcam)^75^. Detergent was removed from the sample by washing twice with 8 M urea in 100 mM Tris-HCl (pH 8) and centrifuged at 15000 rcf for 15 minutes at room temperature. Cysteines were alkylated by addition of 50 mM iodoacetamide for 20 minutes in the dark, washed three times with 8 M urea and three times with 50 mM ammonium bicarbonate. Proteins were subjected to tryptic digestion (1:100 w/w) overnight at 37 °C. Tryptic peptides were collected by centrifugation and washing of the column with 40 µL of 50 mM ammonium bicarbonate and 50 µL of 500 mM sodium chloride. Eluted tryptic peptides were acidified with 1% v/v formic acid and desalted with C_18_ Omix tips (Agilent).

### Analysis of peptides by MRM mass spectrometry

Following MHC-peptide elution, samples were concentrated to a volume < 10 μL using a Labconco Centrivac concentrator at 40 °C, and the volumes equalised to 20 μL in 0.1% formic acid in MilliQ water. Samples were sonicated in a water bath for 10 minutes and centrifuged for 10 minutes at 15,000 rcf and stored in mass spectrometry vials at 4°C for immediate analysis. Peptide abundance was measured as described by Croft, Smith ^40^. Using a SCIEX QTRAP®5500+ mass spectrometer coupled with an on-line Eksigent Ekspert nanoLC 415 (SCIEX, Toronto, Canada). 10 μL of each sample was directly loaded onto a trap column (ChromXP C_18_, 3 μm 120°A, 350 μm × 0.5 mm [SCIEX]) maintained at an isocratic flow of buffer A at a flow rate of 5 μL/min. Peptides were separated using an analytical column (ChromXP C18, 3 μm 120°A, 75 μm× 15 cm [SCIEX]) by increasing linear concentrations of Buffer B at a flow rate of 300 nL/min for 75 min. In MRM mode, unit resolution was used for Q1 and Q3, coupled to an information-dependent acquisition (IDA) criterion triggering an EPI scan (10,000 Da/sec; rolling CE; unit resolution) following any MRM transition exceeding 100 counts and ignoring the triggering MRM transition for 3 seconds thereafter. For analysis of peptides presented *ex vivo* from splenocytes, the same conditions were used except that samples were acquired on a SCIEX QTRAP® 6500+ mass spectrometer.

### Label free quantification of protein expression

Following desalting of tryptic peptides, the equivalent of ∼2 µg of material was analysed on a Q Exactive Plus Orbitrap mass spectrometer (Thermo Scientific) coupled online to an Ultimate 3000 nano-LC system (Thermo Scientific). Samples were loaded on to an Acclaim PepMap100 trap column (100 µm x 2 cm, nanoViper C_18_, 100 Å pore size; Thermo Scientifc) at a flow rate of 15 µL per minute in 2 % v/v acetonitrile 0.1 % v/v formic acid in water. Peptides were eluted across a PepMap100 C_18_ nano column (75 µm x 50 cm, 3 µm, 100 Åpore size; Thermo Scientific) at 250 nL/min with buffer A utilising a gradient of buffer B as follows: hold at 2.5 % for 2 minutes, ramp to 7.5 % for 1 minute, then 40 % buffer over the course of 120 minutes, with a subsequent ramp to 99% over 5 minutes before equilibrating back to 2.5 % over 1 minute and maintaining this for 20 minutes to end the run. The instrument was operated in DDA mode with the following settings: full MS scan with a mass range of 375-1800 m/z, resolution of 70,000 at m/z 200, automatic gain control (AGC) of 3e6 and maximum ion injection time of 50 ms, with dynamic exclusion set to 20 seconds. The top 12 most intense ions selected for HCD fragmentation. A normalised collision energy (NCE) of 27 was used, with maximum injection time of 120 ms, 1.8 m/z isolation window, 17,500 resolution and AGC of 2e5.

Data was analysed by label free quantitation (LFQ) using Maxquant (1.5.2.8) with a combined database of the reference *Mus musculus* proteome (2016, Uniprot) appended with the Vaccinia Virus Western Reserve strain reference proteome. Carbamidomethyl (C) was set as a fixed modification and with acetyl (protein N-term) and oxidation (M) set as variable modifications. MS/MS tolerance was set to 20 ppm, with PSM, Protein and Site FDR set to 1%. Trypsin was used as the enzyme, with a minimum peptide length of 7 and maximum peptide length of 40. “Match between runs” and “Include contaminants” were set to true. The resulting protein output was analysed by Perseus (1.6.0.7). Proteins identified as “only by site”, “reverse” or “contaminants” were removed and LFQ intensities log_10_ transformed. Missing data was subjected to imputation using the default Perseus method.

### Calculation of peptide abundance

The area for each MRM transition was calculated by Skyline Targeted Mass Spec Environment application (MacCoss Lab Software) (MacLean et al., 2010). The area of the transitions for each peptide was added together to calculate the total peak area. For peptides with internal fragment transitions that could not be incorporated into Skyline, quantification was carried out by extracting the peak area under the curve using Peakview (SCIEX).

To estimate the amount of peptide recovered from each spleen, the total peak area, the ratio of the total peak area of the endogenous peptide with the total peak area of the AQUA peptide was obtained. The total amount of endogenous peptide recovered was calculated by multiplying by the amount of AQUA peptide added to the sample (50 fmoles). The equation is summarised below:

Peptide recovered = total peak area(endogenous) / total peak area(AQUA) × Moles(AQUA)

Epitope abundance and relative protein abundance was measured from two independent experiments.

### Statistical Analysis

Average or mean refer to the arithmetic mean to estimate epitope abundance, and geometric mean to compare relative abundance of protein estimated by LFQ.

Imputation strategy for undetected LFQ values: Undetected proteins were assumed to occur due to low abundance. Imputation strategy was modelled on the strategy described by Tyanova, Temu ^76^. For each sample containing both mouse and VACV proteins, data was log-transformed and the median and standard deviation of the log(LFQ) values was calculated for non-null LFQ values. Null LFQ data was imputed by a random distribution centred 1.8 standard deviations below the LFQ detected median, with a distribution of 0.5× standard deviation.

PCA method: PCA for mouse-, VACV-or both mouse and VACV protein relative abundance were derived log-transformed LFQ values from each replicate using the base stats (4.0.3) in R. Variables were centred by not scaled.

Pairwise spearman / Pearson correlations were calculated in Python on log-transformed values.

Non-immunogenic T cell responses were given an arbitrarily low value of 0.01%. For comparisons involving p:MHC-I affinity, individual epitopes were excluded from analysis where measured p:MHC-I affinity was not available.

VACV protein kinetic class (E1.1, E1.2, I, L) for each protein was derived from the following publications based on VACVWR number: ^52^ ^51^ ^16^

## Supporting information

All supplementary figures

## Acknowledgements

We wish to thank the staff of the ANU APF for animal husbandry. This work was supported by: a Project Grant from the NHMRC (Australia) to DCT, AWP and NPC (APP1084283); an NHMRC Senior Research Fellowship and Investigator Grant (APP1104329 and APP2008990) to DCT; an NHMRC Principal Research Fellowship (APP1137739) to AWP; a Viertel Fellowship, ARC Future Fellowship and NHMRC Program Grant (APP1071916) to NLLG.

## Author contributions

Conceptualization, MJW, NPC, AWP, NLLG and DCT; Methodology, NCP, IEAF, AWP and DCT; Investigation, MJW, NPC, YCW, SS, IEAF and EK; Formal Analysis, MJW and NPC; Data Curation, MJW and NPC; Writing – Original Draft, MJW, NPC and DCT; Writing – Review & Editing, MJW, NPC, AWP and DCT; Funding Acquisition, NLLG, AWP and DCT; Supervision, AWP and DCT.

## Supplementary information

Supplementary Figures – suplementary figures 1-12

Table S1. Excel file – Master sheet with summary of all data

Table S2. Excel file – Peptide sequences and MRM transitions

Table S3. Excel file – Mouse and VACV LFQ from infected cell types *in vitro*

Table S4. Excel file – Epitope abundance per replicate from infected cells *in vitro*

Table S5. Excel file – Epitope abundance per replicate from infected mouse spleens

Table S6. Excel file – Peptide-specific CD8^+^ T cell responses to VACV following intravenous infection

## References

1. Burrows, S.R., Rossjohn, J. & McCluskey, J. Have we cut ourselves too short in mapping CTL epitopes? Trends in immunology 27, 11–16 (2006).

2. Pishesha, N., Harmand, T.J. & Ploegh, H.L. A guide to antigen processing and presentation. Nature Reviews Immunology 22, 751–764 (2022).

3. Bennink, J.R., Yewdell, J.W., Smith, G. & Moss, B. Anti-influenza virus cytotoxic T lymphocytes recognize the three viral polymerases and a nonstructural protein: responsiveness to individual viral antigens is major histocompatibility complex controlled. Journal of virology 61, 1098–1102 (1987).

4. Croft, N.P. et al. Most viral peptides displayed by class I MHC on infected cells are immunogenic. Proceedings of the National Academy of Sciences 116, 3112–3117 (2019).

5. Yewdell, J.W. Confronting complexity: real-world immunodominance in antiviral CD8+ T cell responses. Immunity 25, 533–543 (2006).

6. Aki, M. et al. Interferon-γ induces different subunit organizations and functional diversity of proteasomes. The Journal of Biochemistry 115, 257–269 (1994).

7. Glynne, R. et al. A proteasome-related gene between the two ABC transporter loci in the class II region of the human MHC. Nature 353, 357–360 (1991).

8. Hisamatsu, H. et al. Newly identified pair of proteasomal subunits regulated reciprocally by interferon gamma. The Journal of experimental medicine 183, 1807–1816 (1996).

9. Kingsbury, D.J., Griffin, T.A. & Colbert, R.A. Novel propeptide function in 20 S proteasome assembly influences β subunit composition. Journal of Biological Chemistry 275, 24156–24162 (2000).

10. Martinez, C.K. & Monaco, J.J. Homology of proteasome subunits to a major histocompatibility complex-linked LMP gene. Nature 353, 664–667 (1991).

11. Nandi, D., Jiang, H. & Monaco, J.J. Identification of MECL-1 (LMP-10) as the third IFN-gamma-inducible proteasome subunit. Journal of immunology (Baltimore, Md.: 1950) 156, 2361–2364 (1996).

12. de Verteuil, D. et al. Deletion of immunoproteasome subunits imprints on the transcriptome and has a broad impact on peptides presented by major histocompatibility complex I molecules. Molecular & Cellular Proteomics 9, 2034–2047 (2010).

13. Huber, E.M. et al. Immuno-and constitutive proteasome crystal structures reveal differences in substrate and inhibitor specificity. Cell 148, 727–738 (2012).

14. Mishto, M. et al. Proteasome isoforms exhibit only quantitative differences in cleavage and epitope generation. European Journal of Immunology 44, 3508–3521 (2014).

15. Bevan, M.J. Cross-priming for a secondary cytotoxic response to minor H antigens with H-2 congenic cells which do not cross-react in the cytotoxic assay. The Journal of experimental medicine 143, 1283–1288 (1976).

16. Assarsson, E. et al. Kinetic analysis of a complete poxvirus transcriptome reveals an immediate-early class of genes. Proceedings of the National Academy of Sciences 105, 2140–2145 (2008).

17. Apcher, S. et al. Major source of antigenic peptides for the MHC class I pathway is produced during the pioneer round of mRNA translation. Proceedings of the National Academy of Sciences 108, 11572–11577 (2011).

18. Schubert, U. et al. Rapid degradation of a large fraction of newly synthesized proteins by proteasomes. Nature 404, 770–774 (2000).

19. Yewdell, J.W., Antón, L.C. & Bennink, J.R. Defective ribosomal products (DRiPs): a major source of antigenic peptides for MHC class I molecules? Journal of immunology (Baltimore, Md.: 1950) 157, 1823–1826 (1996).

20. Yewdell, J.W. Not such a dismal science: the economics of protein synthesis, folding, degradation and antigen processing. Trends in cell biology 11, 294–297 (2001).

21. Colbert, J.D., Farfán-Arribas, D.J. & Rock, K.L. Substrate-induced protein stabilization reveals a predominant contribution from mature proteins to peptides presented on MHC class I. The Journal of Immunology 191, 5410–5419 (2013).

22. Dolan, B.P. et al. Distinct pathways generate peptides from defective ribosomal products for CD8+ T cell immunosurveillance. The Journal of Immunology 186, 2065–2072 (2011).

23. Hickman, H.D. et al. Toward a definition of self: proteomic evaluation of the class I peptide repertoire. The Journal of Immunology 172, 2944–2952 (2004).

24. Wu, T. et al. Quantification of epitope abundance reveals the effect of direct and cross-presentation on influenza CTL responses. Nature communications 10, 2846 (2019).

25. Dinter, J. et al. Different antigen-processing activities in dendritic cells, macrophages, and monocytes lead to uneven production of HIV epitopes and affect CTL recognition. The Journal of Immunology 193, 4322–4334 (2014).

26. Lazaro, E. et al. Differential HIV epitope processing in monocytes and CD4 T cells affects cytotoxic T lymphocyte recognition. The Journal of infectious diseases 200, 236–243 (2009).

27. Tscharke, D.C., Croft, N.P., Doherty, P.C. & La Gruta, N.L. Sizing up the key determinants of the CD8+ T cell response. Nature Reviews Immunology 15, 705–716 (2015).

28. Kotturi, M.F. et al. Naive precursor frequencies and MHC binding rather than the degree of epitope diversity shape CD8+ T cell immunodominance. J Immunol 181, 2124–2133 (2008).

29. Witney, M.J. & Tscharke, D.C. BMX-A and BMX-S: Accessible cell-free methods to estimate peptide-MHC-I affinity and stability. Mol Immunol 161, 1–10 (2023).

30. Restifo, N.P. et al. Antigen processing in vivo and the elicitation of primary CTL responses. Journal of immunology (Baltimore, Md.: 1950) 154, 4414–4422 (1995).

31. Wong, Y.C. et al. Modified vaccinia virus Ankara can induce optimal CD8+ T cell responses to directly primed antigens depending on vaccine design. Journal of Virology 93, e01154–01119 (2019).

32. Chen, W., Antón, L.C., Bennink, J.R. & Yewdell, J.W. Dissecting the multifactorial causes of immunodominance in class I–restricted T cell responses to viruses. Immunity 12, 83–93 (2000).

33. Crotzer, V.L. et al. Immunodominance among EBV-derived epitopes restricted by HLA-B27 does not correlate with epitope abundance in EBV-transformed B-lymphoblastoid cell lines. The Journal of Immunology 164, 6120–6129 (2000).

34. Gallimore, A., Dumrese, T., Hengartner, H., Zinkernagel, R.M. & Rammensee, H.-G. Protective immunity does not correlate with the hierarchy of virus-specific cytotoxic T cell responses to naturally processed peptides. The Journal of experimental medicine 187, 1647-b–1657 (1998).

35. Levitsky, V., Zhang, Q.-J., Levitskaya, J. & Masucci, M.G. The life span of major histocompatibility complex-peptide complexes influences the efficiency of presentation and immunogenicity of two class I-restricted cytotoxic T lymphocyte epitopes in the Epstein-Barr virus nuclear antigen 4. The Journal of experimental medicine 183, 915–926 (1996).

36. Pamer, E.G. Direct sequence identification and kinetic analysis of an MHC class I-restricted Listeria monocytogenes CTL epitope. Journal of immunology (Baltimore, Md.: 1950) 152, 686–694 (1994).

37. Tsomides, T.J. et al. Naturally processed viral peptides recognized by cytotoxic T lymphocytes on cells chronically infected by human immunodeficiency virus type 1. The Journal of experimental medicine 180, 1283–1293 (1994).

38. van Els, C.A. et al. A single naturally processed measles virus peptide fully dominates the HLA-A* 0201-associated peptide display and is mutated at its anchor position in persistent viral strains. European journal of immunology 30, 1172–1181 (2000).

39. Vijh, S. & Pamer, E.G. Immunodominant and subdominant CTL responses to Listeria monocytogenes infection. Journal of immunology (Baltimore, Md.: 1950) 158, 3366–3371 (1997).

40. Croft, N.P. et al. Kinetics of antigen expression and epitope presentation during virus infection. PLoS pathogens 9, e1003129 (2013).

41. Upton, C., Slack, S., Hunter, A.L., Ehlers, A. & Roper, R.L. Poxvirus orthologous clusters: toward defining the minimum essential poxvirus genome. Journal of virology 77, 7590–7600 (2003).

42. Briody, B.A. Response of mice to ectromelia and vaccinia viruses. Bacteriol Rev 23, 61–95 (1959).

43. Goulding, J., Bogue, R., Tahiliani, V., Croft, M. & Salek-Ardakani, S. CD8 T cells are essential for recovery from a respiratory vaccinia virus infection. J Immunol 189, 2432–2440 (2012).

44. Hersperger, A.R., Siciliano, N.A., DeHaven, B.C., Snook, A.E. & Eisenlohr, L.C. Epithelial immunization induces polyfunctional CD8+ T cells and optimal mousepox protection. J Virol 88, 9472–9475 (2014).

45. Jiang, X. et al. Skin infection generates non-migratory memory CD8+ T(RM) cells providing global skin immunity. Nature 483, 227–231 (2012).

46. Lin, L.C., Flesch, I.E. & Tscharke, D.C. Immunodomination during peripheral vaccinia virus infection. PLoS pathogens 9, e1003329 (2013).

47. Xu, R.-H., Remakus, S., Ma, X., Roscoe, F. & Sigal, L.J. Direct presentation is sufficient for an efficient anti-viral CD8+ T cell response. PLoS pathogens 6, e1000768 (2010).

48. Xu, R., Johnson, A.J., Liggitt, D. & Bevan, M.J. Cellular and humoral immunity against vaccinia virus infection of mice. J Immunol 172, 6265–6271 (2004).

49. Shen, Z., Reznikoff, G., Dranoff, G. & Rock, K.L. Cloned dendritic cells can present exogenous antigens on both MHC class I and class II molecules. Journal of immunology (Baltimore, Md.: 1950) 158, 2723–2730 (1997).

50. Trinchieri, G., Aden, D.P. & Knowles, B.B. Cell-mediated cytotoxicity to SV40-specific tumour-associated antigens. Nature 261, 312–314 (1976).

51. Yang, Z. et al. Expression profiling of the intermediate and late stages of poxvirus replication. Journal of virology 85, 9899–9908 (2011).

52. Yang, Z., Bruno, D.P., Martens, C.A., Porcella, S.F. & Moss, B. Simultaneous high-resolution analysis of vaccinia virus and host cell transcriptomes by deep RNA sequencing. Proceedings of the National Academy of Sciences 107, 11513–11518 (2010).

53. Norbury, C.C. et al. CD8+ T cell cross-priming via transfer of proteasome substrates. Science 304, 1318–1321 (2004).

54. W. Yewdell, J. The seven dirty little secrets of major histocompatibility complex class I antigen processing. Immunological reviews 207, 8–18 (2005).

55. Koutsakos, M. et al. Human CD8+ T cell cross-reactivity across influenza A, B and C viruses. Nature immunology 20, 613–625 (2019).

56. Nicholas, B. et al. Immunopeptidomic analysis of influenza A virus infected human tissues identifies internal proteins as a rich source of HLA ligands. PLoS Pathogens 18, e1009894 (2022).

57. Goncalves, G. et al. IFNγ modulates the immunopeptidome of triple negative breast cancer cells by enhancing and diversifying antigen processing and presentation. Frontiers in Immunology 12, 645770 (2021).

58. Kincaid, E.Z. et al. Mice completely lacking immunoproteasomes show major changes in antigen presentation. Nature immunology 13, 129–135 (2012).

59. Abelin, J.G. et al. Mass Spectrometry Profiling of HLA-Associated Peptidomes in Mono-allelic Cells Enables More Accurate Epitope Prediction. Immunity 46, 315–326 (2017).

60. Bassani-Sternberg, M., Pletscher-Frankild, S., Jensen, L.J. & Mann, M. Mass spectrometry of human leukocyte antigen class I peptidomes reveals strong effects of protein abundance and turnover on antigen presentation. Mol Cell Proteomics 14, 658–673 (2015).

61. Fortier, M.H. et al. The MHC class I peptide repertoire is molded by the transcriptome. J Exp Med 205, 595–610 (2008).

62. Hoof, I., van Baarle, D., Hildebrand, W.H. & Kesmir, C. Proteome sampling by the HLA class I antigen processing pathway. PLoS Comput Biol 8, e1002517 (2012).

63. Zanker, D., Waithman, J., Yewdell, J.W. & Chen, W. Mixed proteasomes function to increase viral peptide diversity and broaden antiviral CD8+ T cell responses. The Journal of Immunology 191, 52–59 (2013).

64. Flesch, I.E. et al. Altered CD8+ T cell immunodominance after vaccinia virus infection and the naive repertoire in inbred and F1 mice. The journal of immunology 184, 45–55 (2010).

65. La Gruta, N.L. et al. A virus-specific CD8+ T cell immunodominance hierarchy determined by antigen dose and precursor frequencies. Proceedings of the National Academy of Sciences 103, 994–999 (2006).

66. Nayar, R. et al. Graded levels of IRF4 regulate CD8+ T cell differentiation and expansion, but not attrition, in response to acute virus infection. The Journal of immunology 192, 5881–5893 (2014).

67. Obar, J.J., Khanna, K.M. & Lefrançois, L. Endogenous naive CD8+ T cell precursor frequency regulates primary and memory responses to infection. Immunity 28, 859–869 (2008).

68. Norbury, C.C., Malide, D., Gibbs, J.S., Bennink, J.R. & Yewdell, J.W. Visualizing priming of virus-specific CD8+ T cells by infected dendritic cells in vivo. Nature immunology 3, 265–271 (2002).

69. Hickman, H.D. et al. Direct priming of antiviral CD8+ T cells in the peripheral interfollicular region of lymph nodes. Nature immunology 9, 155–165 (2008).

70. Serna, A., Ramirez, M.C., Soukhanova, A. & Sigal, L.J. Cutting edge: efficient MHC class I cross-presentation during early vaccinia infection requires the transfer of proteasomal intermediates between antigen donor and presenting cells. The Journal of Immunology 171, 5668–5672 (2003).

71. Lev, A. et al. The exception that reinforces the rule: crosspriming by cytosolic peptides that escape degradation. Immunity 28, 787–798 (2008).

72. Madaan, A., Verma, R., Singh, A.T., Jain, S.K. & Jaggi, M. A stepwise procedure for isolation of murine bone marrow and generation of dendritic cells. J Biol Methods 1, e1 (2014).

73. Flesch, I.E., Hollett, N.A., Wong, Y.C. & Tscharke, D.C. Linear fidelity in quantification of anti-viral CD8+ T cells. PLoS One 7, e39533 (2012).

74. Purcell, A.W., Ramarathinam, S.H. & Ternette, N. Mass spectrometry–based identification of MHC-bound peptides for immunopeptidomics. Nature protocols 14, 1687–1707 (2019).

75. Wiśniewski, J.R., Zougman, A., Nagaraj, N. & Mann, M. Universal sample preparation method for proteome analysis. Nature methods 6, 359–362 (2009).

76. Tyanova, S. et al. The Perseus computational platform for comprehensive analysis of (prote) omics data. Nature methods 13, 731–740 (2016).

