## Supplementary material for "A quantitative approach to poxvirus infection reveals general correlates of antigen presentation and immunogenicity for viral CD8^+^ T cell epitopes": All supplementary figures

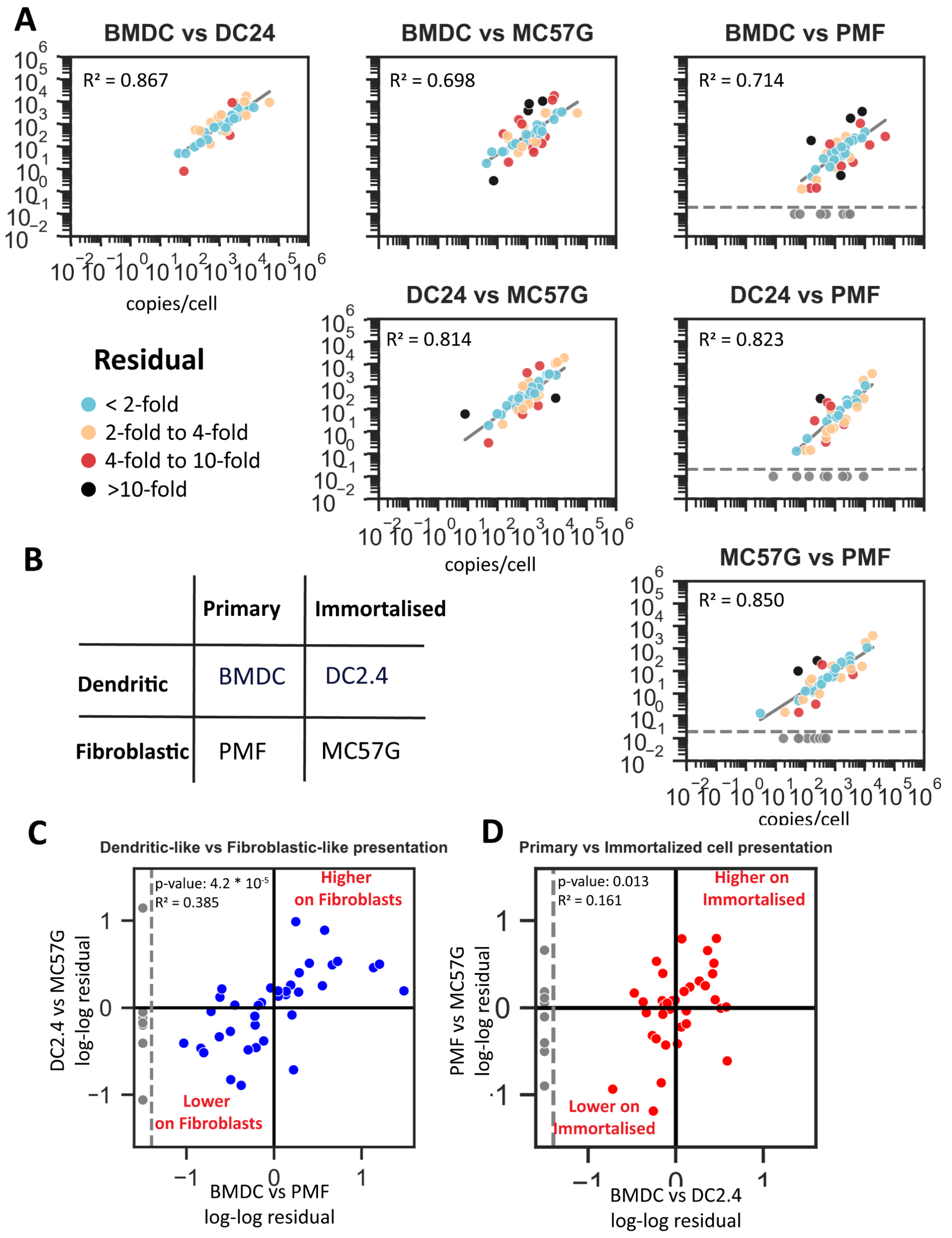


**Supplementary Figure 1: Pairwise comparisons of VACV epitopes abundance on different cell types.** BMDC, DC2.4, MC57G and PMF cells were infected with VACV. p:MHC-I abundance of 45 epitopes were measured by MRM at 0.5, 2.5, 4.5, 6.5 and 8.5h post infection in two independent experiments. The sum of the average abundance for each epitope at all timepoints were compared between different cell lines. A) Pairwise correlation of p:MHC-I abundance measured on cell types. Statistics calculated using Pearson correlation from log-converted values. Diagonal lines represent line of best fit. Points are colour coded by distance from the line of best fit as indicated. Undetected p:MHC-I are shown below the horizontal dashed line but excluded from statistical analysis. B) Summary of shared features between BMDC, DC2.4, MC57G and PMF cells. C-D) Correlation of residuals from pairwise comparisons of BMDC v PMF and DC2.4 v MC57G (C) or BMDC v DC2.4 and PMF v MC57G (D).

**A**

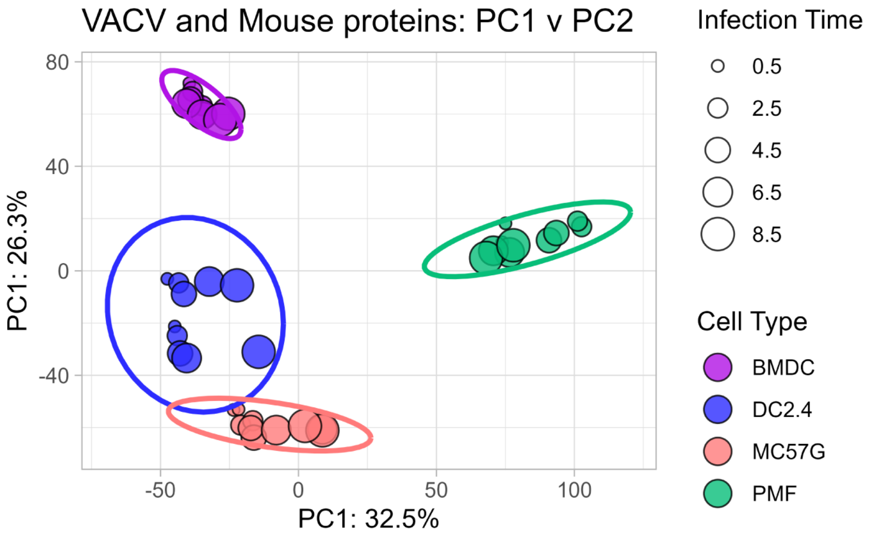


**B**

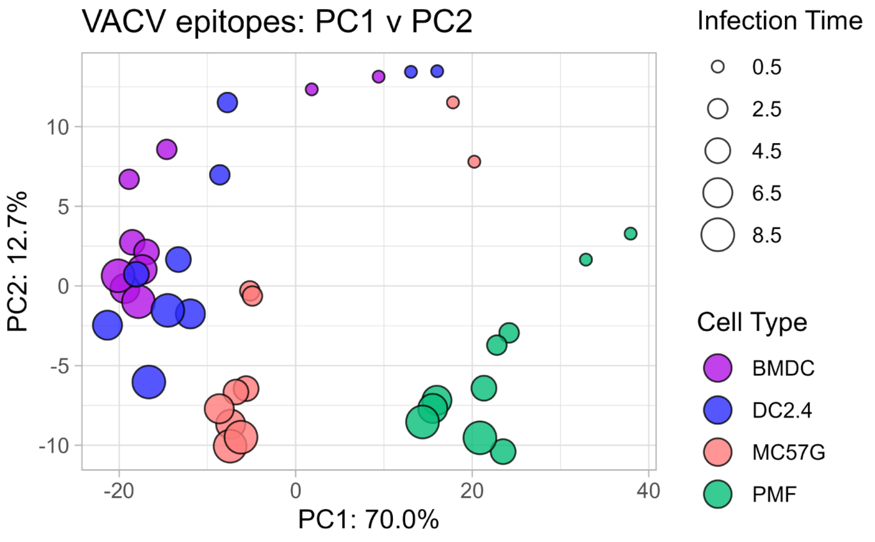


**Supplementary Figure 2: PCA plots for complete protein and epitope datasets.**
PCA plots describing PC1 and PC2 axis for combined mouse and VACV protein LFQ intensity (A) or VACV p:MHC-I abundance (B) for each replicate and timepoint on VACV-infected BMDC, DC2.4, MC57G and PMF cell types. Colours indicate cell type, the size of each point identifies the time after infection.

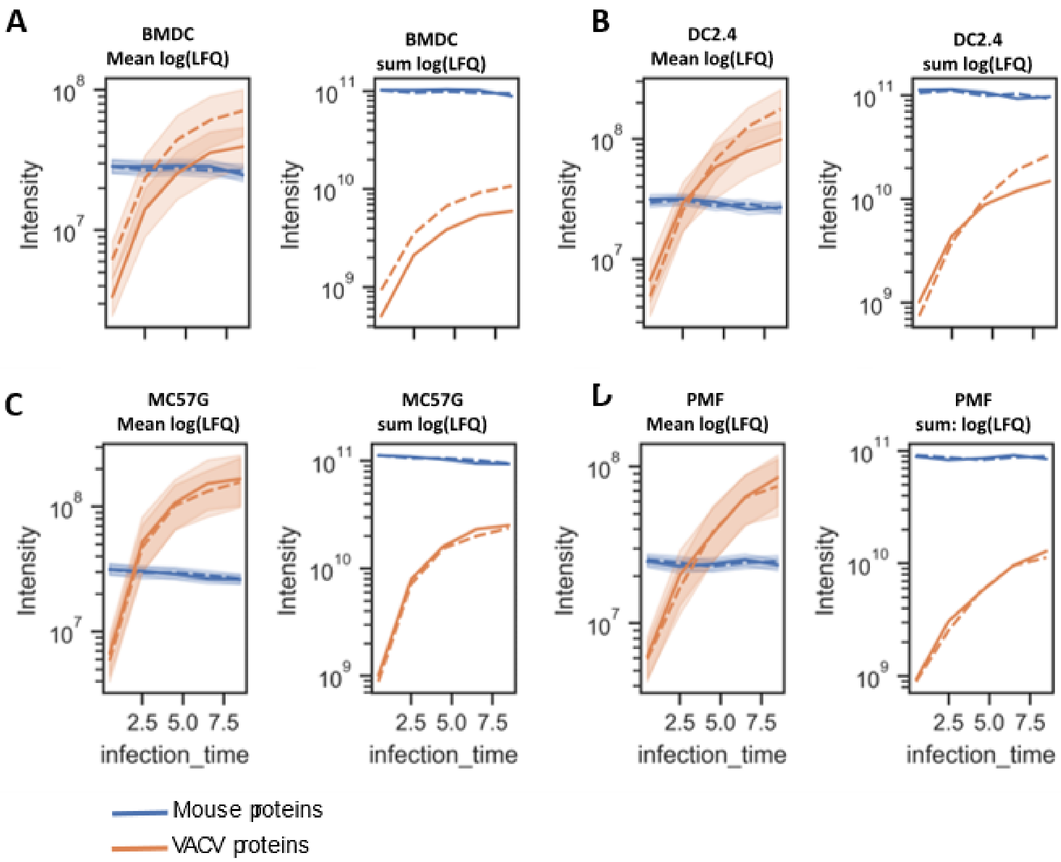


**Supplementary Figure 3: Relative protein abundances of VACV and mouse proteins over time.**
Each plot compares the geometric mean or the sum of the relative abundance of mouse (blue) and VACV (orange) proteins on BMDC (A), DC2.4 (B), MC57G (C) and PMF cells (D) over time. Each line represents a separate replicate. Shaded areas represent the variation in abundance of individual proteins.

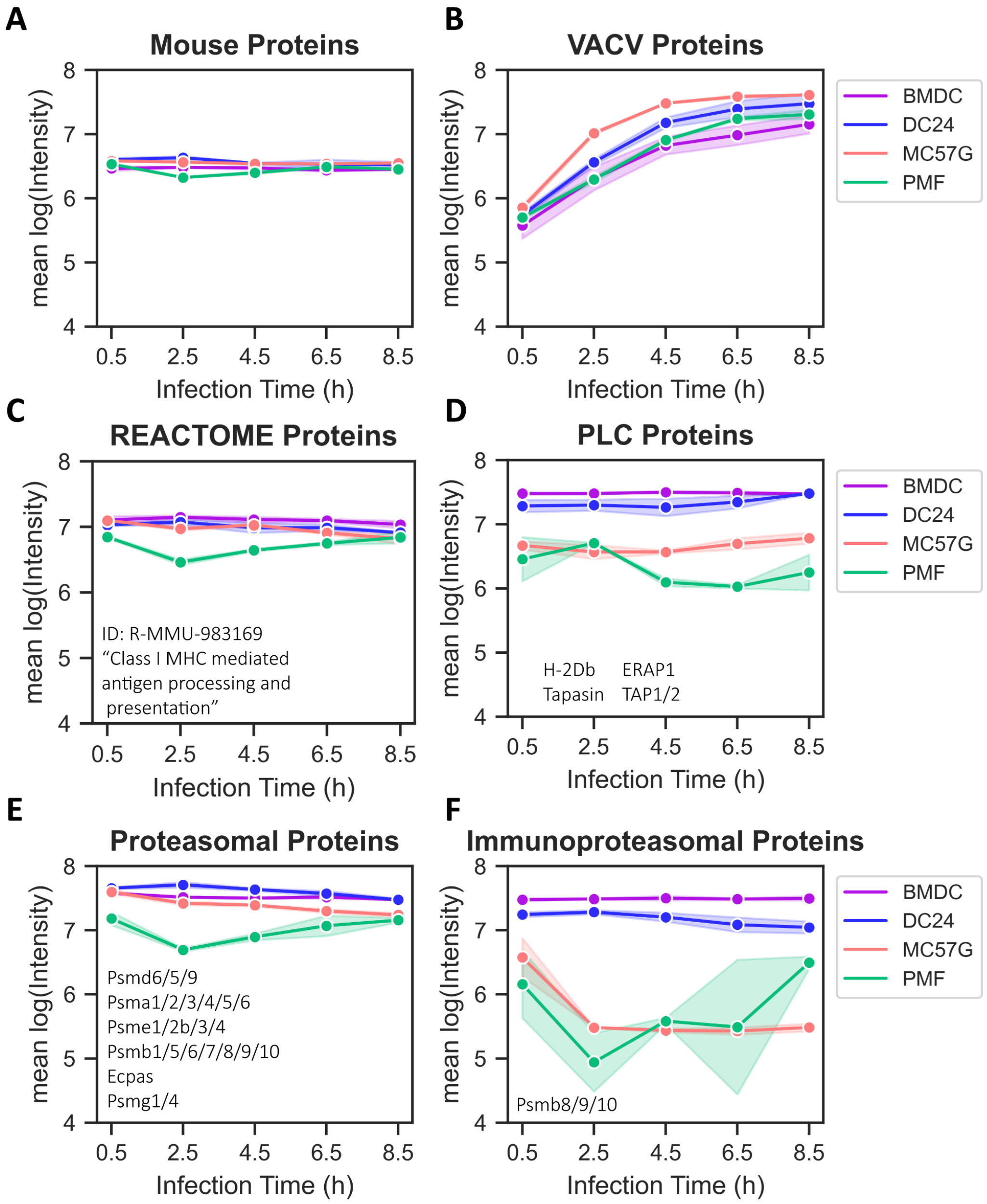


**Supplementary Figure 4: Abundance of Mouse proteins associated with p:MHC-I processing and presentation.** The geometric mean of proteins at each time point and cell type for subsets of proteins that represented all mouse proteins (A), all VACV proteins (B), matched REACTOME pathway proteins associated with MHC-I processing (ID: R-MMU:983169) (C), proteins associated with the peptide loading complex (PLC) (D), proteasomal degradation (E) or immunoproteasome subunits (F). Gene names in the plot identify the respective protein included in the analysis.

**A**

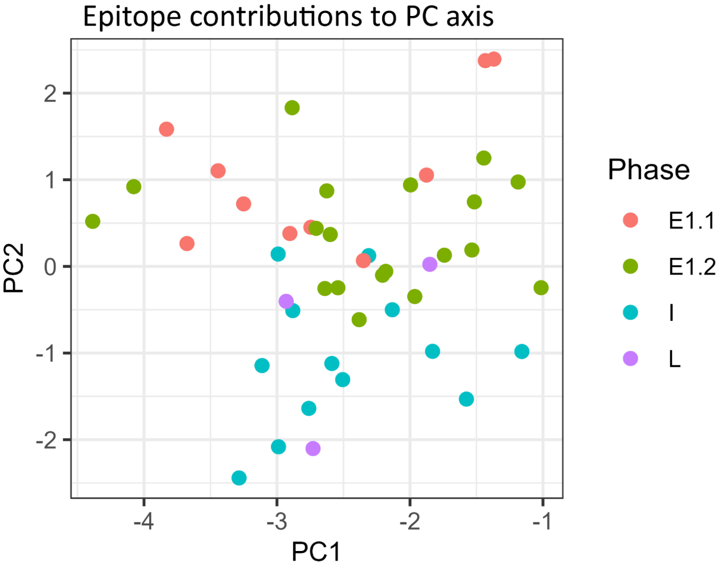


**B**

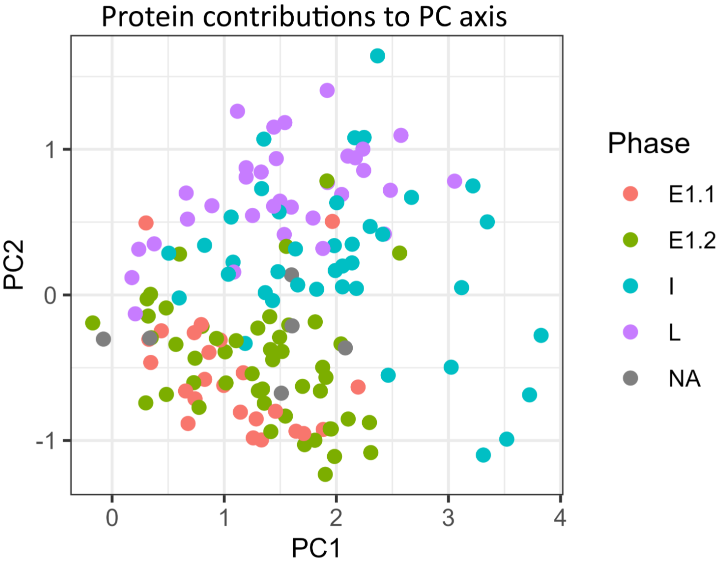


**Supplementary Figure 5: Variable contributions towards PC1 and PC2 axes.**
The relative contributions of variables towards PC1 and PC2 axes from the PCA analysis of either VACV epitopes (A) or VACV proteins (B) as shown in Fig. 1 and Fig. 2 respectively. Each point identifies the loading of each epitope or protein contributing to PC1 and PC2, multiplied by the size of the PC eigenvalue. Points are coloured by the kinetic class (E1.1, E1.2, Intermediate (I), Late (L)) of the respective VACV protein.

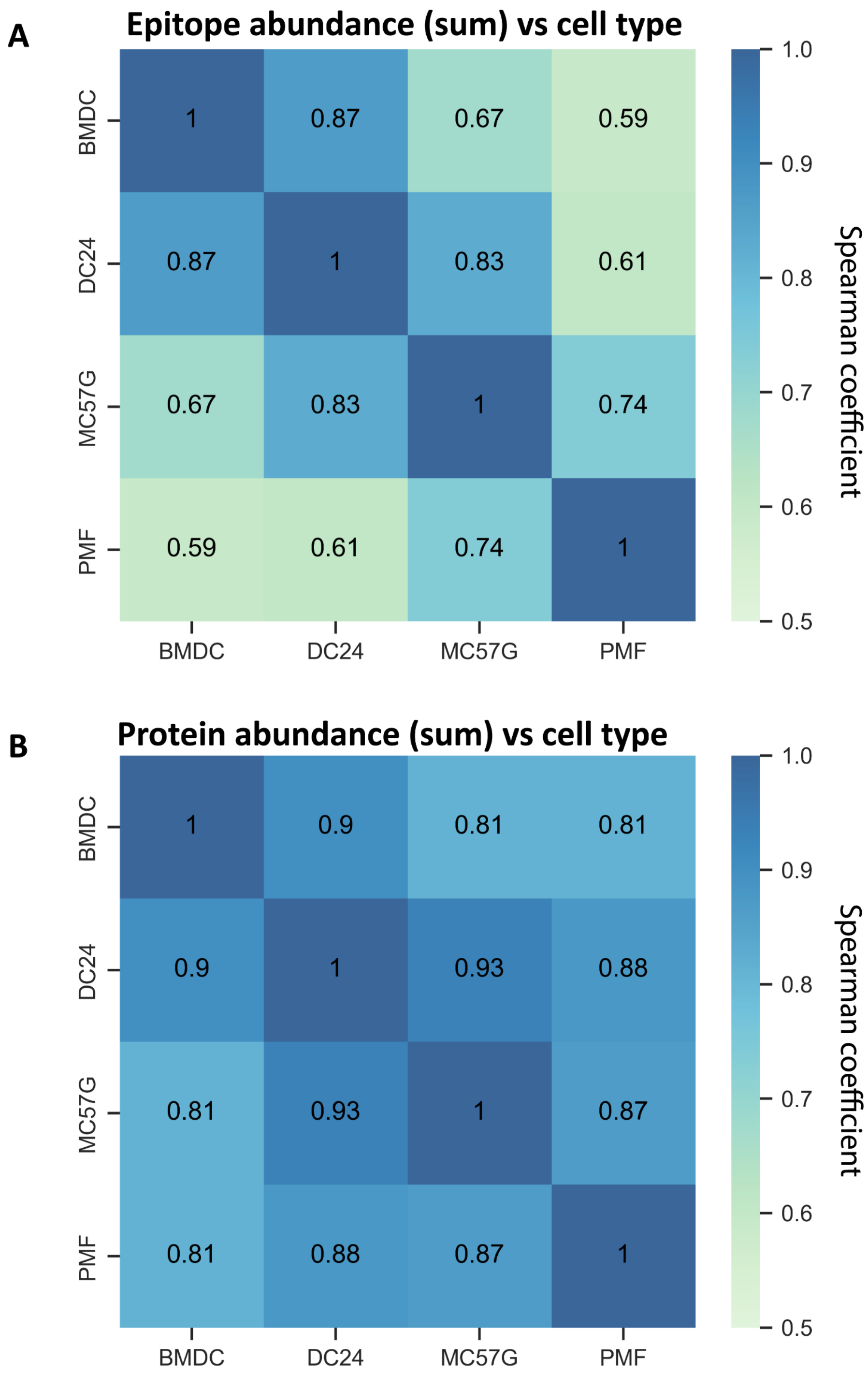


**Supplementary Figure 6: Conservation of p:MHC-I kinetic class between cell types.**
Spearman correlation coefficients for pairwise comparisons between epitope (A) and protein (B) abundance measured on each cell type. The abundance of each epitope and protein was evaluated as the averaged sum of values across all timepoints on each cell type.

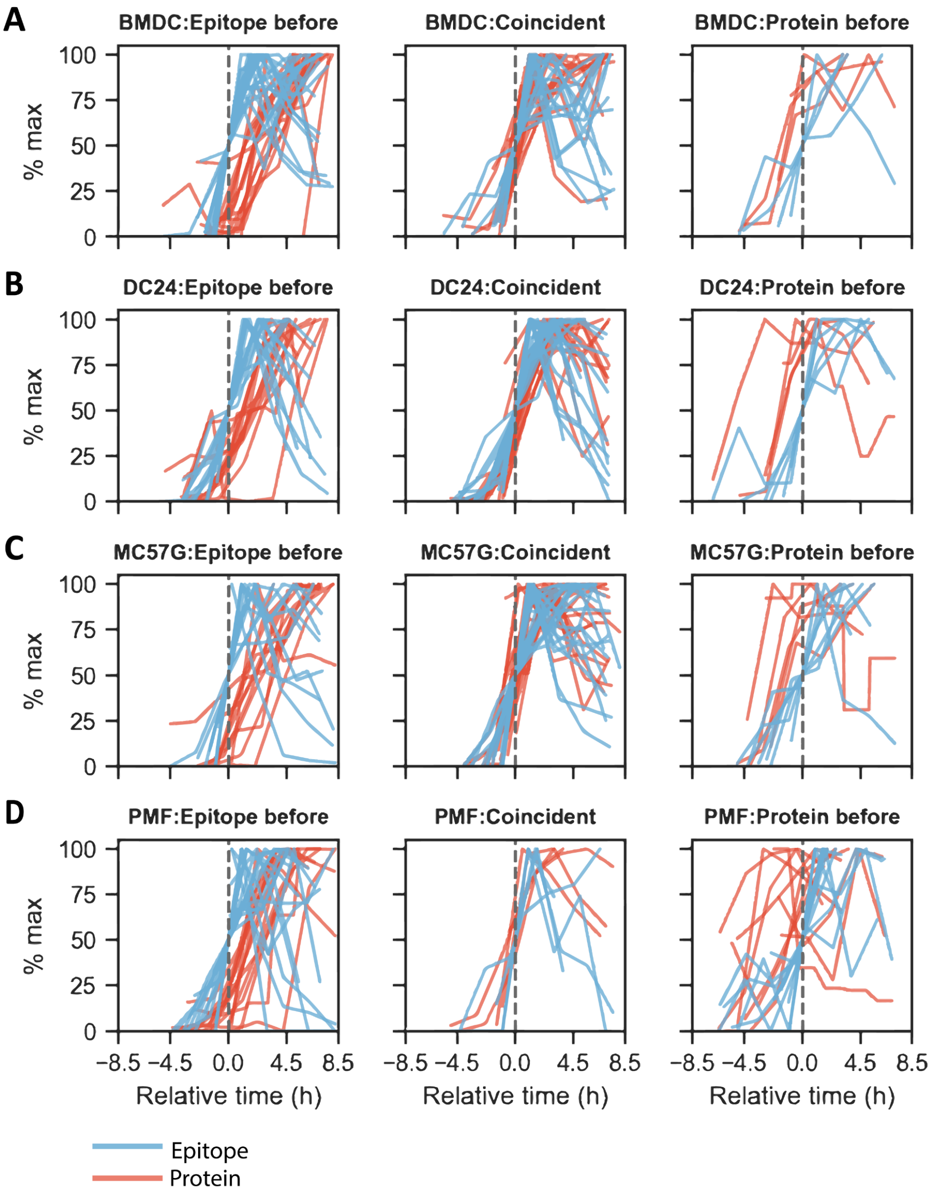


**Supplementary Figure 7: VACV epitope presentation kinetics, classified as before, coincident or after protein expression.**

The average p:MHC-I abundance (MRM MS) and VACV protein abundance (LFQ) from two replicates were normalised with respect to the maximum value over the time-course. Each epitope was matched to the corresponding VACV antigen. The time to 50% of the maximum value ($t_{max/2}$) for epitope and protein abundance was calculated on VACV-infected BMDC (A), DC2.4 (B), MC57G (C) and PMF cells (D). Epitope presentation before, coincident with, or after protein expression was classified according to whether the epitope $t_{max/2}$ was at least 1 hour earlier, less than one hour, or more than one hour after the $t_{max/2}$ of the respective protein. Relative time represents protein and epitope abundance translated along the x-axis such that epitope $t_{max/2}$ is centred at 0. Blue lines and red lines identify average kinetics of each epitope and protein respectively.

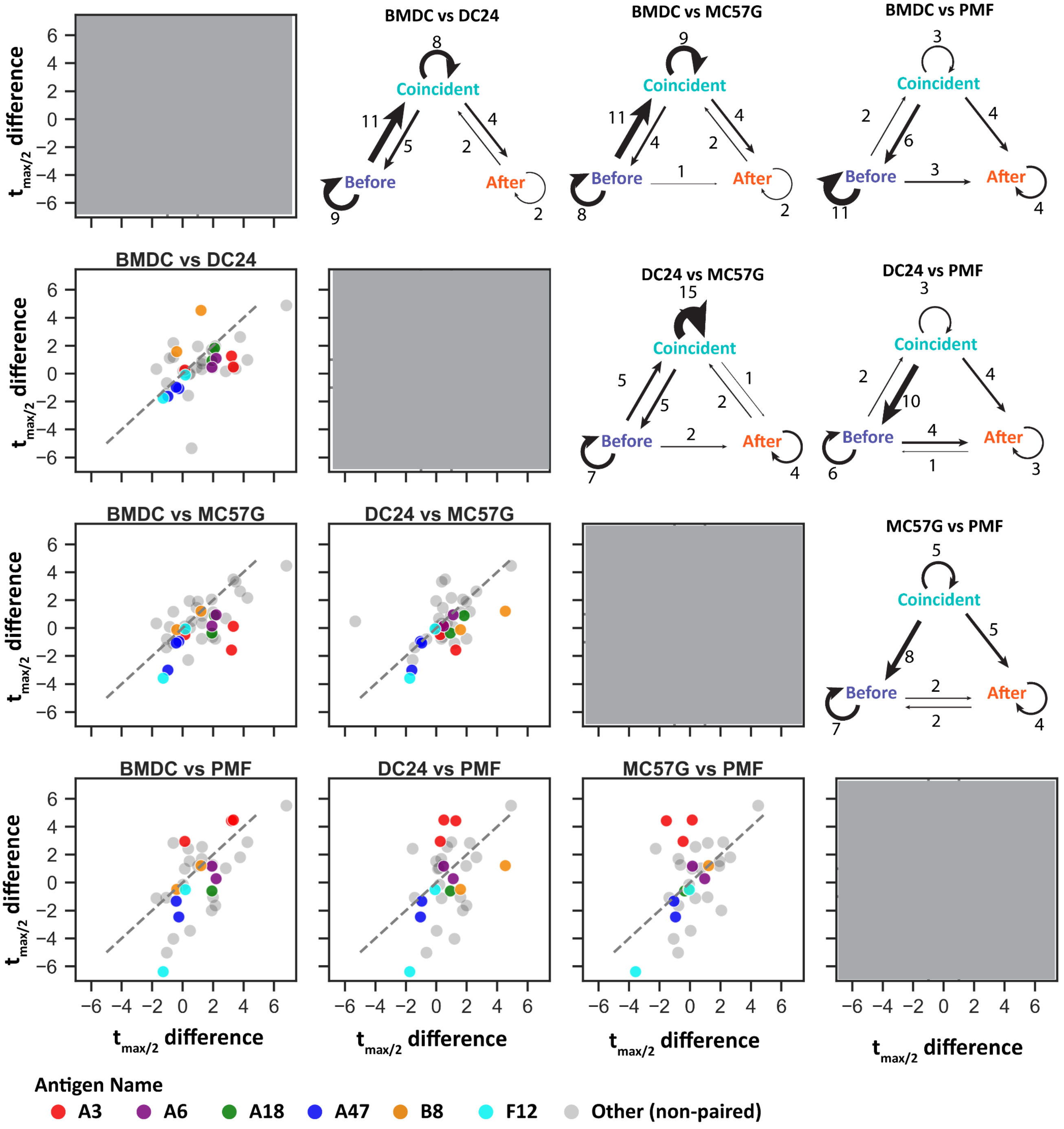


**Supplementary Figure 8: Conservation of p:MHC-I kinetic class between cell types.**
Upper right: Arrows indicate the number of p:MHC-I that were redefined into new classification between the two respective cell types. Size of arrows are scaled to the number indicated.
Lower left: Pairwise comparisons describing the difference between the $t_{max/2}$ of the protein and epitope on the respective cell types. p:MHC-I derived from the same VACV protein are coloured similarly as indicated. Unrelated p:MHC-I shown in grey. The diagonal dashed line represents position expected for p:MHC-I with the same time gap between epitope and protein $t_{max/2}$ for the two cell types.

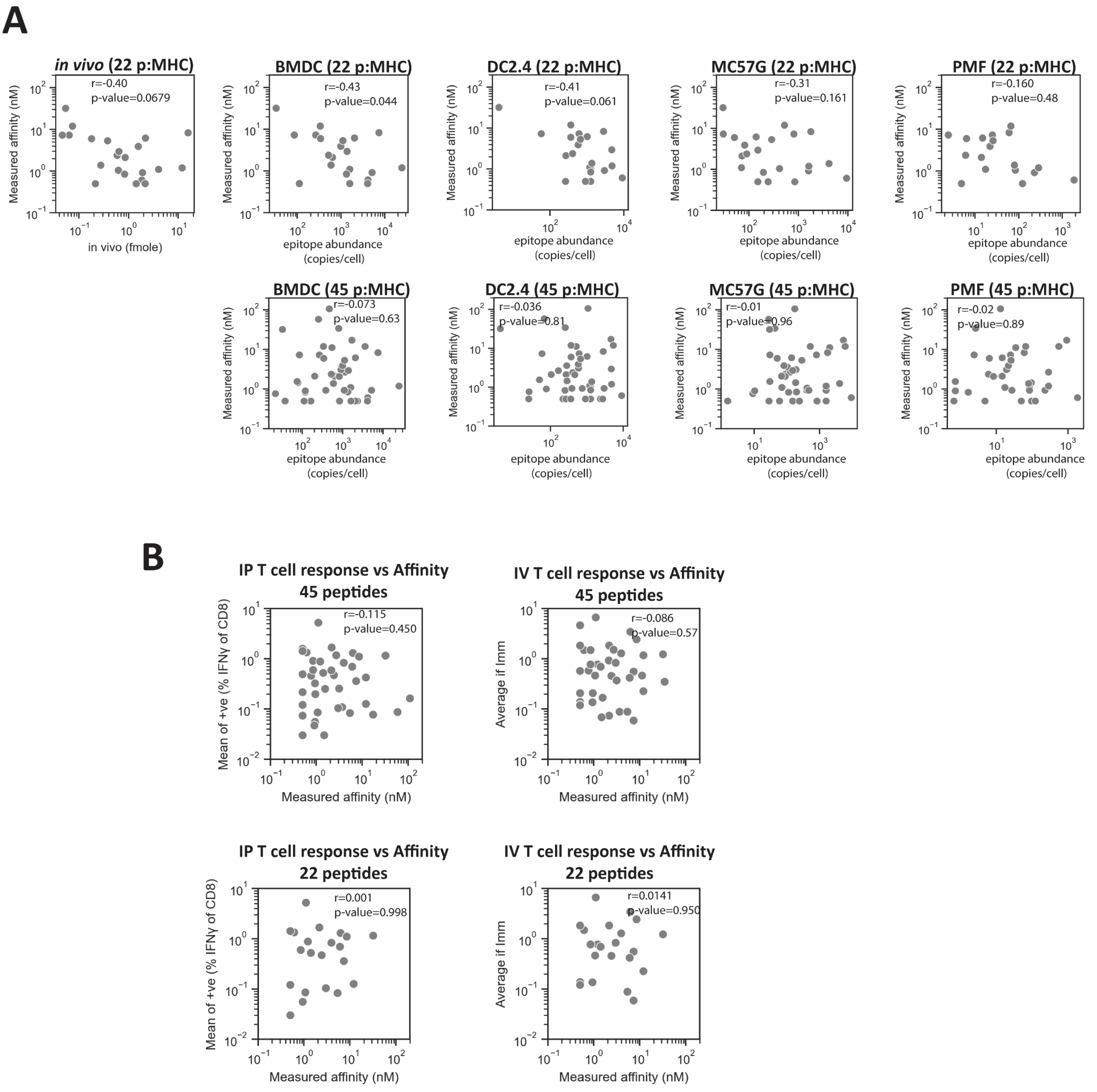


**Supplementary Figure 9:**

A) Pairwise comparison of epitope abundance and measure affinity for the 22 epitopes measured *in vivo* or abundance of the 45 p:MHC-I measured on cell types *in vitro*. Statistic: spearman correlation.

B) Spearman correlation of p:MHC-I affinity and mean T cell response during VACV infection following intraperitoneal infection (Croft et al., 2019) and intravenous infection (current publication).

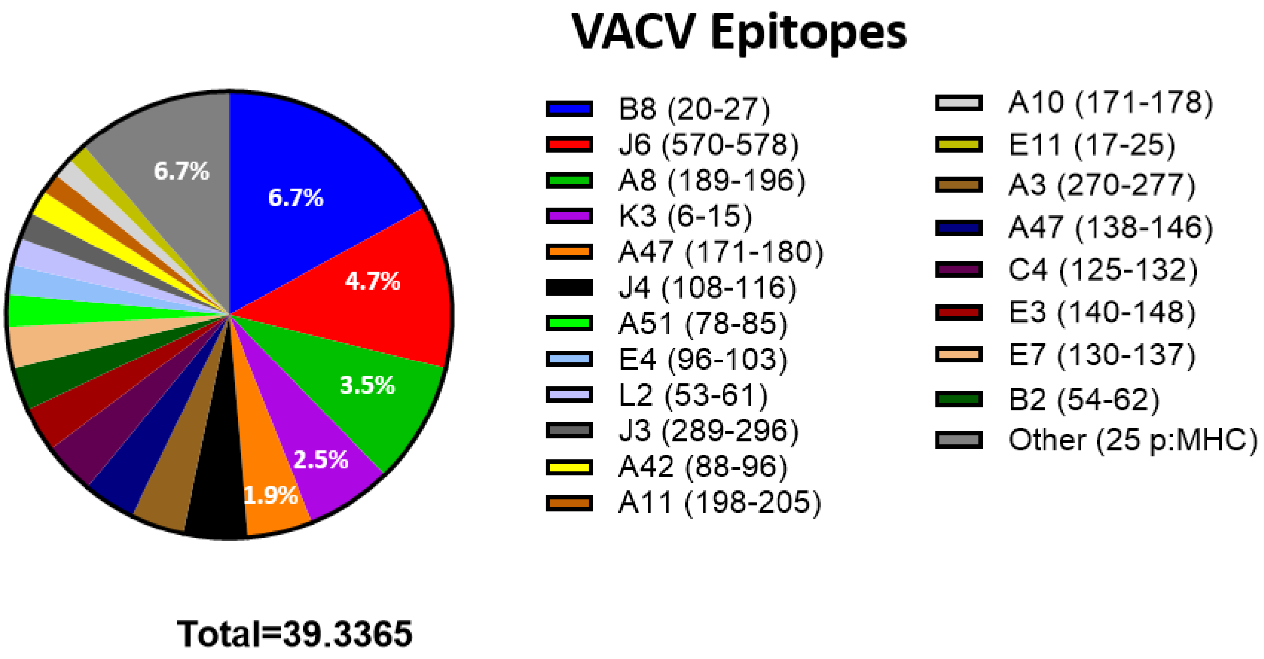


**Supplementary Figure 10: Visualizing the share of the total IFNγ+ T cell response attributed to each p:MHCI.** Each peptide-specific T cell response is represented by a coloured fraction of the total IFNγ+ T cell response that could be accounted for by the 45 p:MHC-I investigated in this study.

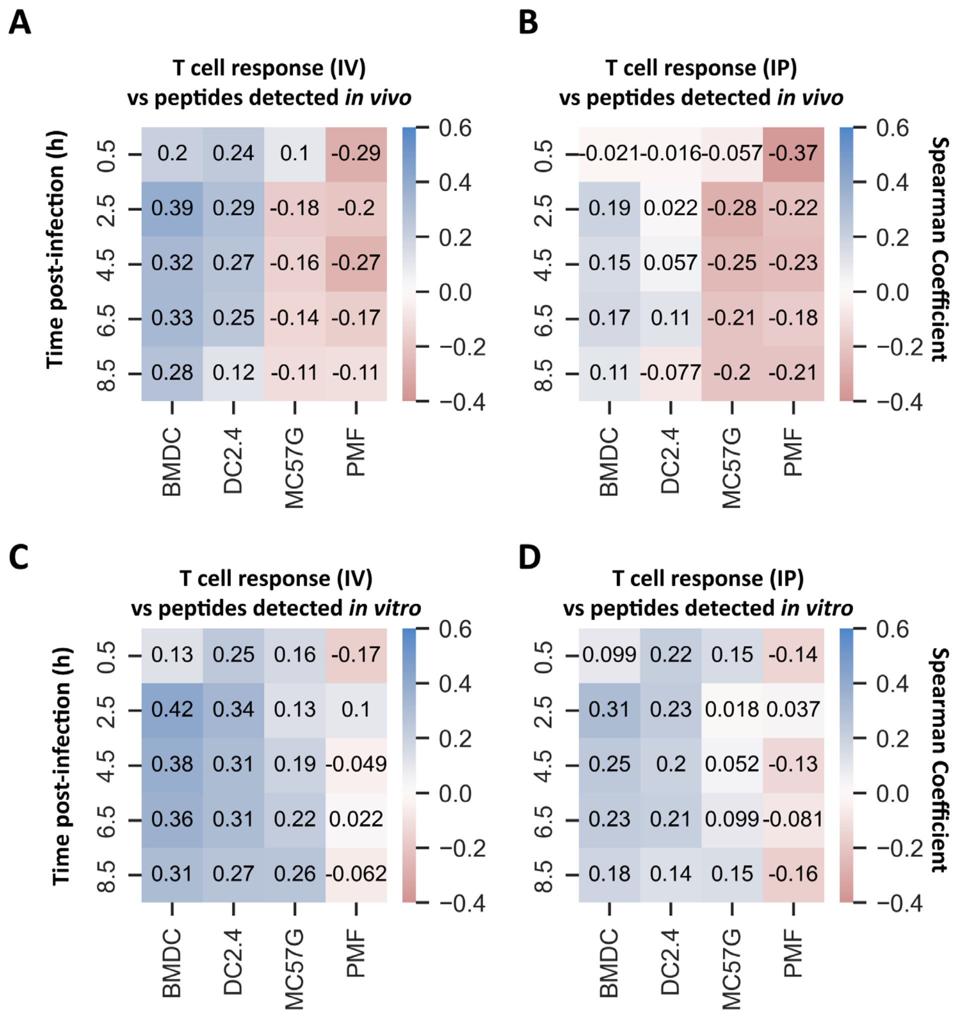


**Supplementary Figure 11: Exhaustive pairwise comparisons of p:MHC-I abundance and T cell responses.** The spearman correlation coefficients comparing epitope abundance on each cell type and time-point post infection with peptide-specific T cell responses following intravenous (A,C) or intraperitoneal (B,D) infection and in C57BL/6 mice. Comparisons were made either restricted to the set of peptides detected *in vivo* (A,B) or the complete set of 45 epitopes (C,D). Epitope abundances not detected in a given timepoint were given an arbitrarily low value of 0.01 copies/cell.

### Supp Fig 12 A) BMDC

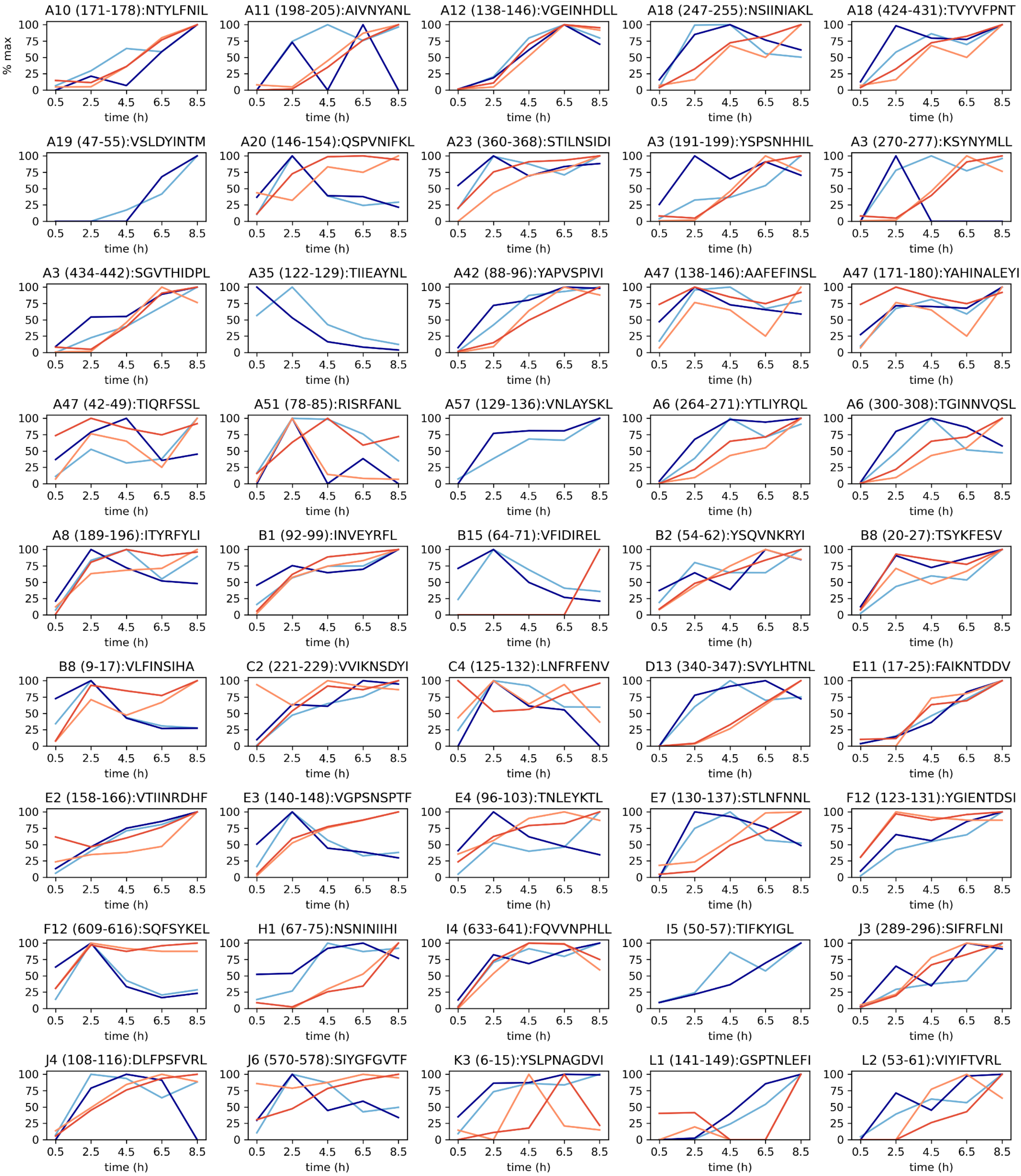


**Supplementary Figure 12: Paired kinetics of p:MHC-I and the source antigen on *in vitro* infected cells.** p:MHC-I and protein abundance was normalized to the maximum value for each replicate measured on BMDCs (A), DC2.4 (B), MC57G (C) or PMF (D) cells. p:MHC-I were matched with the relevant VACV source protein. Each line represents a single replicate for p:MHC-I (dark blue, light blue) or protein (red, orange) abundance. Data for p:MHC-I are annotated by position within the respective VACV protein and sequence.

**Supp Fig 12 B) DC2.4**

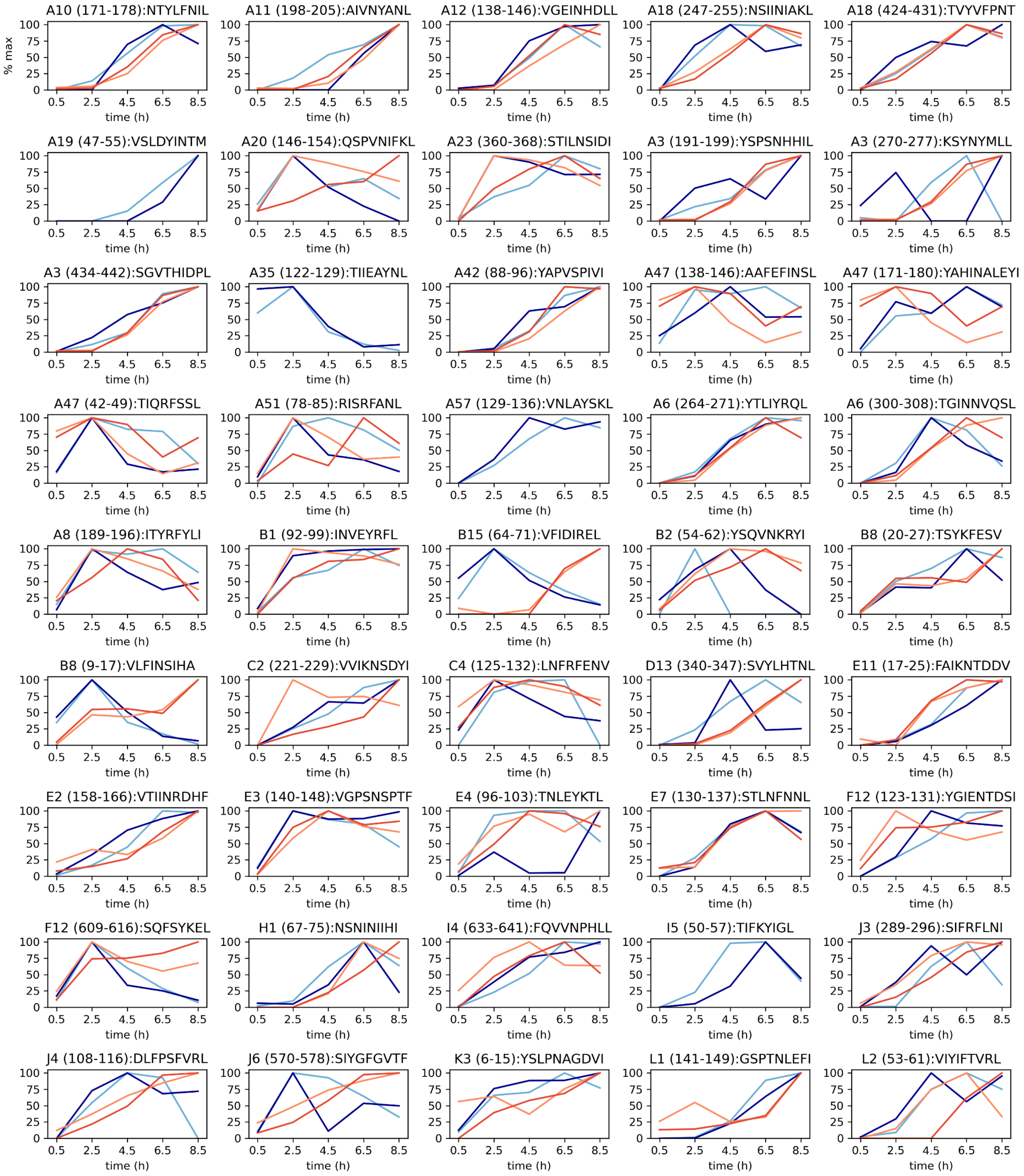


**Supplementary Figure 12: Paired kinetics of p:MHC-I and the source antigen on *in vitro* infected cells.** p:MHC-I and protein abundance was normalized to the maximum value for each replicate measured on BMDCs (A), DC2.4 (B), MC57G (C) or PMF (D) cells. p:MHC-I were matched with the relevant VACV source protein. Each line represents a single replicate for p:MHC-I (dark blue, light blue) or protein (red, orange) abundance. Data for p:MHC-I are annotated by position within the respective VACV protein and sequence.

### Supp Fig 12 C) MC57G

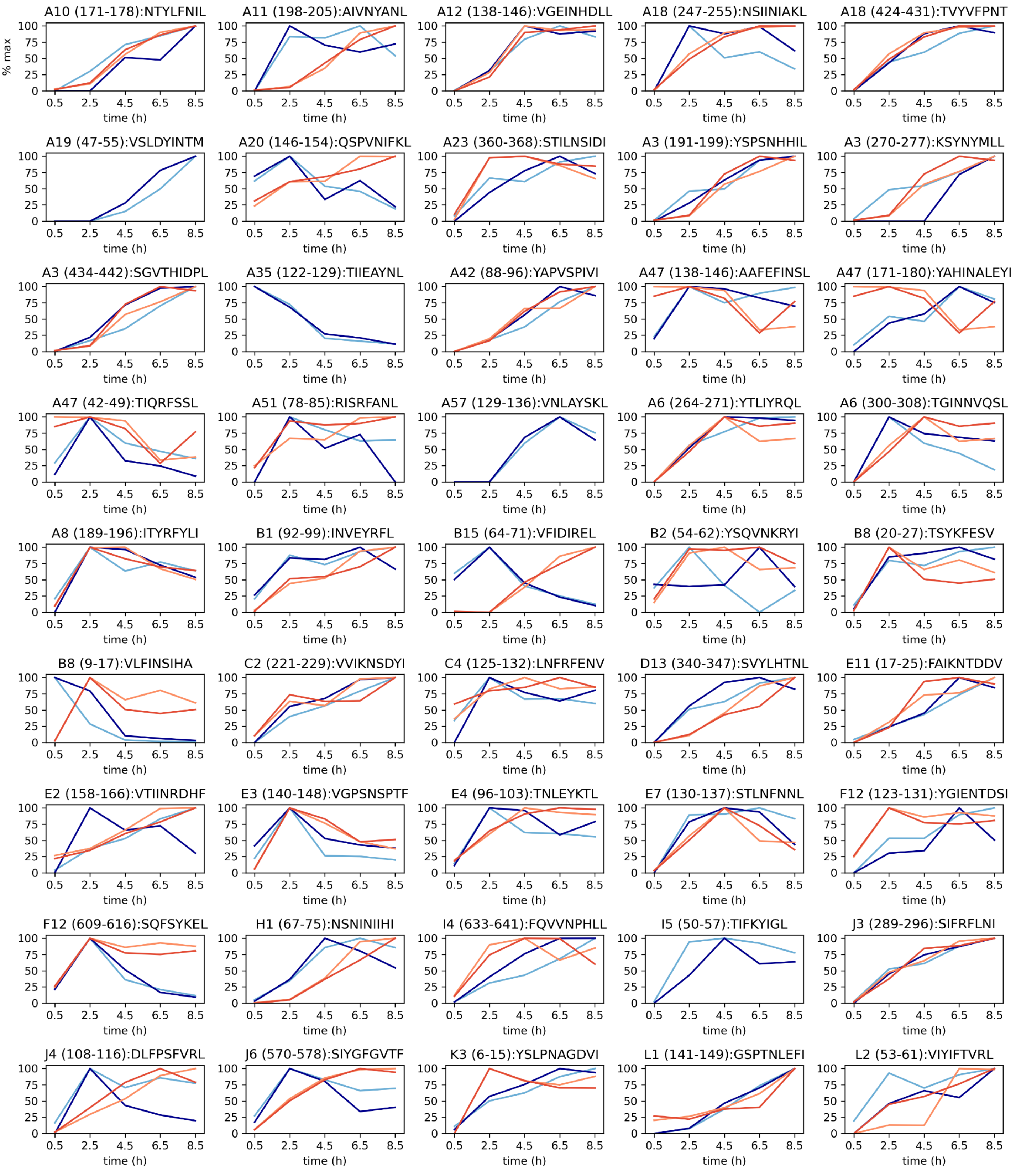


**Supplementary Figure 12: Paired kinetics of p:MHC-I and the source antigen on *in vitro* infected cells.** p:MHC-I and protein abundance was normalized to the maximum value for each replicate measured on BMDCs (A), DC2.4 (B), MC57G (C) or PMF (D) cells. p:MHC-I were matched with the relevant VACV source protein. Each line represents a single replicate for p:MHC-I (dark blue, light blue) or protein (red, orange) abundance. Data for p:MHC-I are annotated by position within the respective VACV protein and sequence.

**Supp Fig 12 D) PMF**

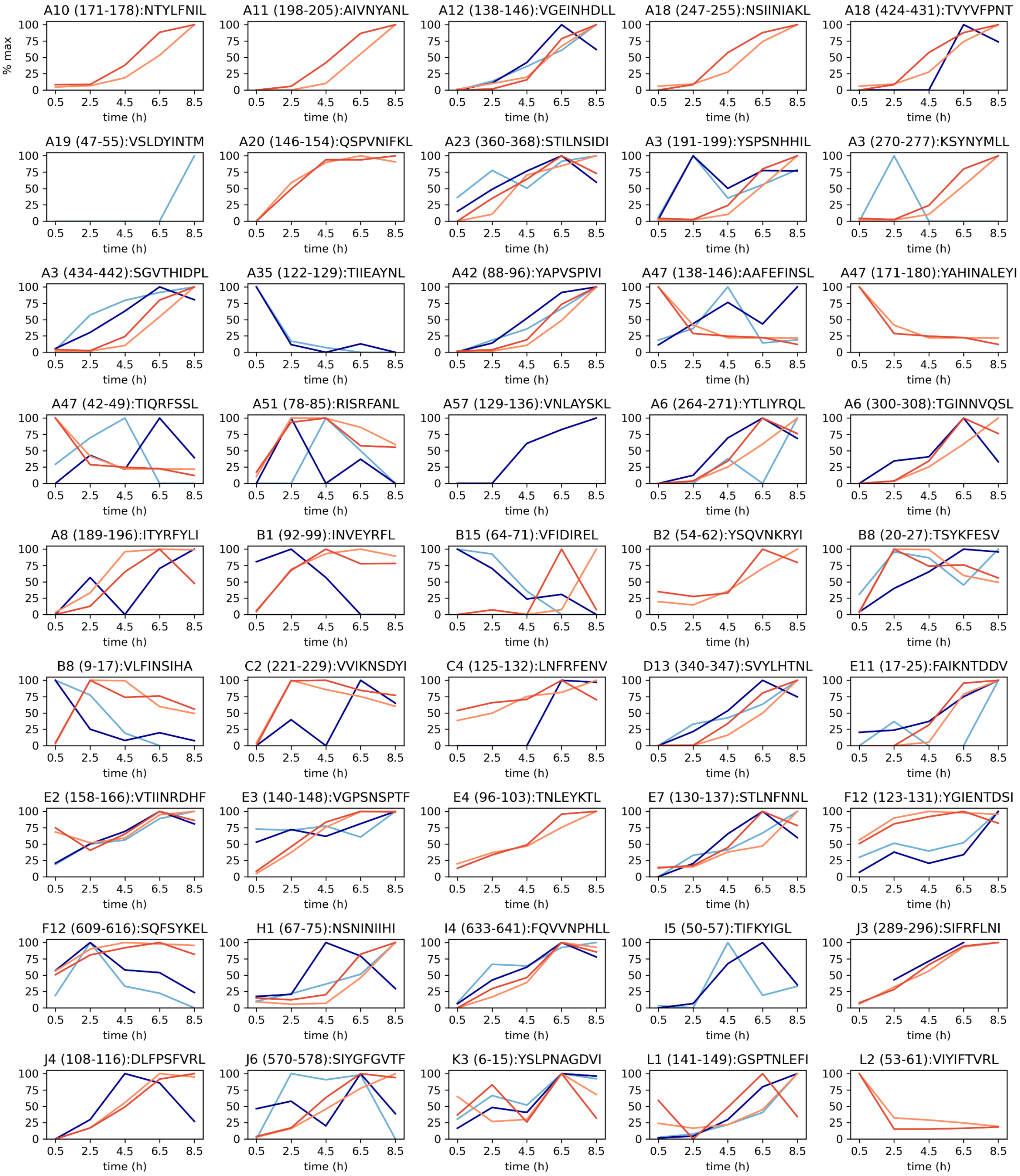


**Supplementary Figure 12: Paired kinetics of p:MHC-I and the source antigen on *in vitro* infected cells.** p:MHC-I and protein abundance was normalized to the maximum value for each replicate measured on BMDCs (A), DC2.4 (B), MC57G (C) or PMF (D) cells. p:MHC-I were matched with the relevant VACV source protein. Each line represents a single replicate for p:MHC-I (dark blue, light blue) or protein (red, orange) abundance. Data for p:MHC-I are annotated by position within the respective VACV protein and sequence.
